# Tree sequences as a general-purpose tool for population genetic inference

**DOI:** 10.1101/2024.02.20.581288

**Authors:** Logan S. Whitehouse, Dylan Ray, Daniel R. Schrider

## Abstract

As population genetics data increases in size new methods have been developed to store genetic information in efficient ways, such as tree sequences. These data structures are computationally and storage efficient, but are not interchangeable with existing data structures used for many population genetic inference methodologies such as the use of convolutional neural networks (CNNs) applied to population genetic alignments. To better utilize these new data structures we propose and implement a graph convolutional network (GCN) to directly learn from tree sequence topology and node data, allowing for the use of neural network applications without an intermediate step of converting tree sequences to population genetic alignment format. We then compare our approach to standard CNN approaches on a set of previously defined benchmarking tasks including recombination rate estimation, positive selection detection, introgression detection, and demographic model parameter inference. We show that tree sequences can be directly learned from using a GCN approach and can be used to perform well on these common population genetics inference tasks with accuracies roughly matching or even exceeding that of a CNN-based method. As tree sequences become more widely used in population genetics research we foresee developments and optimizations of this work to provide a foundation for population genetics inference moving forward.

## INTRODUCTION

Recent decades have seen dramatic growth in the volume, scale, and diversity of population genetic datasets. This explosion in the amount of data has coincided with the development of an expansive set of statistical methods designed to make more accurate and detailed evolutionary inferences from this glut of data. Initial methods often relied on examining the values of statistics that summarize relevant aspects of genetic diversity (Watterson 1975; Nei and Li 1979; Tajima 1989; Fu and Li 1993). For example, statistics quantifying information about the presence of high-frequency derived alleles may be used to identify recent positive selection, which produces an excess of high-frequency derived alleles via genetic hitchhiking (Fay and Wu 2000). In addition to statistics that summarize the distribution of allele frequencies in a population, researchers have designed statistics to capture other properties of genetic variation as well, such as the density of polymorphism (Nei and Li 1979; Nei and Tajima 1981), patterns of haplotypic diversity (Hudson et al. 1994; Sabeti et al. 2002; Voight et al. 2006; Garud et al. 2015), and the extent of linkage disequilibrium (Kelly 1997; Kim and Nielsen 2004). Because such summary statistics necessarily result in the loss of potentially valuable information by representing population genomic diversity by a single number, more recent approaches, such as approximate Bayesian computation (ABC) sought to combine multiple summaries into a higher-dimensional representation of the data that could then be used for inference (Tavare et al. 1997; Pritchard et al. 1999; Beaumont et al. 2002). ABC methods work by comparing large sets of simulations, each summarized by a vector of statistics, to identify the evolutionary model and/or parameter combination that best matches the observed data. Importantly, the summary statistic vector can capture a variety of properties of genetic variation, that together may be informative about the evolutionary models/parameters being considered.

More recently, researchers have begun developing supervised machine learning (SML) methods for population genetic inference (Schrider and Kern 2018; Korfmann, Gaggiotti, et al. 2023). Unlike traditional applications of supervised learning, these approaches use simulated data to train and test methods before applying them to empirical datasets. This reliance on simulated training data makes SML approaches to population genetic inference similar in principle to ABC, although these methods often have greater accuracy and computational efficiency (Pudlo et al. 2016; Raynal et al. 2019). SML methods have been applied to a variety of population genetic tasks including detecting positive selection (Pavlidis et al. 2010; Lin et al. 2011; Ronen et al. 2013; Pybus et al. 2015; Kern and Schrider 2018; Mughal and DeGiorgio 2019; Mughal et al. 2020; Arnab et al. 2022; Hejase et al. 2022; Whitehouse and Schrider 2022), demographic parameter estimation (Sheehan and Song 2016; Wang et al. 2021) and model inference (Sanchez et al. 2021), inferring recombination rates (Gao et al. 2016; Adrion et al. 2020), detecting introgression (Schrider et al. 2018; Gower et al. 2021; Ray et al. 2023), and numerous other applications (Battey et al. 2020; Battey et al. 2021; Yelmen et al. 2021; Booker et al. 2023; Smith et al. 2023). The initial applications of SML to population genetic problems took as their input a vector of summary statistics, and yielded sizable accuracy gains on several tasks (reviewed in (Schrider and Kern 2018)). Because representing genetic diversity within a population by a set of statistics may still result in valuable information being lost, researchers have more recently begun to experiment with deep learning methods that are capable of directly examining a population genetic alignment as input and learning their own set of features to extract from these data (Chan, Perrone, Spence, et al. 2018; Flagel et al. 2019). Although deep learning has allowed for a multitude of significant advances in population genetics (reviewed in (Korfmann, Gaggiotti, et al. 2023)), one important drawback of this approach is that population genetic alignments are an inefficient format for representing data (Kelleher et al. 2019).

Because of the large amount of information that must be stored in onboard GPU memory, examining large genomic regions or entire chromosomes and/or large sample sizes is a significant computational challenge for deep learning methods that take alignments as their input. One approach to alleviate this issue is to store genetic information as the sequence of marginal trees along the chromosome describing the shared evolutionary history of the sequenced genomes (Kelleher et al. 2016), with adjacent genealogies in this sequence separated by the breakpoints of recombination events that resulted in a change to the sample’s genealogy (Wiuf and Hein 1999). Concise representations of this sequence of genealogies (“succinct tree sequences”) can be used to efficiently store all of the information present in a population genetic alignment, and this innovation has led to dramatic improvements in simulation efficiency (Kelleher et al. 2018; Haller and Messer 2022). Relatedly, this data representation can be used as a hyper-efficient form of lossless compression for population genetic data (Kelleher et al. 2019; Wong et al. 2024), and can in principle describe the complete evolutionary history of a sample. These advantages have motivated the development of methods to use sequence data to infer the sequence of marginal genealogies (Speidel et al. 2019; Zhang et al. 2023) or even the full ancestral recombination graph (ARGs; Rasmussen *et al*. 2014; Kelleher *et al*. 2019; Mahmoudi *et al*. 2022).

In addition to inferring genealogies from population genetic data, there is growing interest in conducting inference on the genealogies themselves. For example, succinct tree sequences have been used to efficiently compute commonly used summary statistics with dramatic improvements in speed and memory usage (Ralph et al. 2020). Because these genealogies record a sample’s evolutionary history, an exciting prospect is to directly examine them to make downstream inferences (Brandt et al. 2024; Lewanski et al. 2024), and several methods have been devised to this end (Stern et al. 2019; Fan et al. 2023). Researchers have also begun developing methods using ARGs or summaries thereof as input to machine learning methods for evolutionary inference (Hejase et al. 2022; Korfmann, Sellinger, et al. 2023; Pearson and Durbin 2023). Given that when applied to real datasets such methods will be run on genealogies that are inferred with a considerable amount of error (Zhang et al. 2023) one concern may be that ARG-based inference methods using erroneous input information may produce less satisfactory results than methods directly acting on the input sequence data. However, even in situations where genealogical inference may introduce error, SML methods that are trained to handle potential errors/biases in their input genealogies could potentially circumvent this problem (Mo and Siepel 2023), resulting in successful use of SML on inferred genealogies.

With these potential strengths and weaknesses of genealogy-based inference in mind, we sought to assess whether deep learning approaches trained on the sequence of marginal genealogies are likely to add value in comparison to methods acting directly on population genetic alignments. We address this question by assessing the performance of graph convolutional networks (GCNs), a generalization of convolutional neural networks (CNNs) to handle graphical input data. We train GCNs to solve common tasks in population genetic inference: detecting loci that have experienced introgression between closely related species, identifying selective sweeps, estimating recombination rates, and inferring demographic parameters (the same tasks examined previously by (Flagel et al. 2019)). Encouragingly, we find that GCNs trained to use inferred genealogies yield roughly equivalent or even better accuracy to those trained on population genetic alignments. We close with a discussion of the implications of our GCN’s performance for future population genetic research leveraging this increasingly commonly used representation of evolutionary histories and genetic variation data.

## RESULTS

Our goal is to assess the general effectiveness of the approach of performing population genetic inference from inferred tree sequences. To this end, we evaluate the performance of graph convolutional networks (GCNs), a type of deep neural network that takes graphical data as its input, when applied to tree sequences inferred by *Relate*. First, a note on terminology: we note that *Relate* does not attempt to infer which internal nodes in different trees correspond to one another (even though this sequence of trees will be autocorrelated as expected), although the complete ARG would contain this information (Brandt et al. 2024; Lewanski et al. 2024). Our goal was to design a neural network that could use such data as its input. Therefore, we refer to the sequence of marginal trees examined by our GCN as a “tree sequence” rather than an ARG. However, we note that the “tree sequences” used as input to our GCN should not be confused with “succinct tree sequences”, a data structure that records ARG information in a highly efficient manner (Kelleher et al. 2016; Kelleher et al. 2019; Wong et al. 2024) but that we do not use here.

We gauge the success of GCNs by comparing their performance to that of CNNs trained on image representations of the input population genetic alignment data—this comparison is an informative measuring stick given that the CNN-image approach has been shown to be highly effective on a number of population genetic problems, as discussed above. We focus our assessment on the four tasks examined by Flagel *et al*. (Flagel et al. 2019): estimating historical recombination rates, detecting and classifying selective sweeps, detecting the presence and directionality of introgression, and inferring demographic parameters. We begin with an overview of GCNs and how they differ from CNNs as well as their application to tree sequences before addressing the two approaches’ performance on each problem in turn.

### Graph convolutional networks on tree sequences

Convolutional neural networks (CNNs) have proved remarkably effective in a variety of fields (Hochreiter and Schmidhuber 1997; LeCun et al. 1998; Erhan et al. 2013) (Gu et al. 2018; Li et al. 2022), and in recent years have been successfully applied to a number of problems in population genetics (Sheehan and Song 2016; Schrider and Kern 2018; Korfmann, Gaggiotti, et al. 2023). These networks work by extracting features from images, or other structured data that can be represented as tensors of two or more dimensions such as audio or video data, using convolutional filters. By using a series of convolution filter layers, often combined with pooling layers that downsample the data, CNNs are able to extract features within the data that are pertinent to the target task. These features are typically then passed as input into a one or more fully connected layers that then emit the final prediction: a vector of values corresponding to either to class membership probabilities in the case of classification (e.g. with each class representing a different possible object present in the image), or to estimated parameter/attribute values in the case of regression.

The convolutional filters in a CNN, each of which is smaller than the input to which they are applied, work by striding across the input image and at each step performing a transformation of the portion of the input image that is currently “covered” by the filter (LeCun et al. 1989). The transformed value is a function of the pixels in the input image and also the values of the weights of the filter (specifically, the grand sum of the Hadamard product between the filter and the corresponding portion of the input). The filter then moves to another location of the input and repeats the process, until a filtered version of the image has been produced. During training, the weights of the convolutional filters are adjusted to minimize error on the task at hand, thereby producing a set of filters that are capable of extracting low-level structural features (e.g. edges in an image) from the input image that are informative for downstream classification/regression by the rest of the neural network. When multiple layers of convolutions are stacked on top of each other, with the output from one layer passed to the next, features about higher level structures present in the input (e.g. shapes in the image) can be extracted.

GCNs work similarly to CNNs in that they extract features from an input using parameterized filters to convolve over the input space. As the name implies however, rather than an image or “image-like” data structure GCN input consists of a graph. This graph is represented as a pair of matrices: 1) an adjacency matrix describing the structure and connectedness of the nodes in the graph, and 2) a feature matrix containing a vector of attributes for each node in the graph. When centered on a focal node, convolutions in GCNs incorporate information from each of its neighbors, with iterative convolutions passing information from further nodes through information sharing and, in the case of graph attention networks (GATs), attention. GCNs are thus a generalization of CNNs, with the latter being a special case where each pixel in the input corresponds to a node within a lattice graph structure. The design of GCNs allows them to learn from explicit connectedness in a graph to make inferences based on topology and features.

Although a relatively new form of neural networks, GCNs have been shown to have excellent accuracy on tasks where graph data is relevant and have become a widely-researched type of neural network (Zhou et al. 2020), however they have had only limited use in population genetics applications (Korfmann, Gaggiotti, et al. 2023; Korfmann, Sellinger, et al. 2023).

Because genealogies are a type of graph, the trees in a tree sequence are a fitting input for a GCN. However, in order to extract the most information possible for inference the spatial relationship between trees must be considered as well. To accomplish this we built a hybrid GCN-RNN (Figure 1) architecture to learn both the graph structure of each tree in a sequence as well as the spatial relationship of trees along a chromosome. Each tree is passed through six graph convolution layers which extract information from an individual tree in the sequence comprising of node features (node age, number of mutations, population label; see Methods for more details) and the connectedness of the graph through adjacency matrix (represented by edge indices).After graph convolution, we pass the node features through two RNNs (recurrent neural networks), the first of which compresses the features from each tree into a single feature vector and the second compresses the features from each sequence of trees into a single vector. This vector is then used as input to a fully connected network (MLP) that is used to optimize output for a given task. This combination of models optimizing for a single task results in an architecture able to learn both the spatial relatedness of nodes within a tree and their relevance as well as the larger-scale structured information of the tree sequence as a whole.

**Figure 1.**
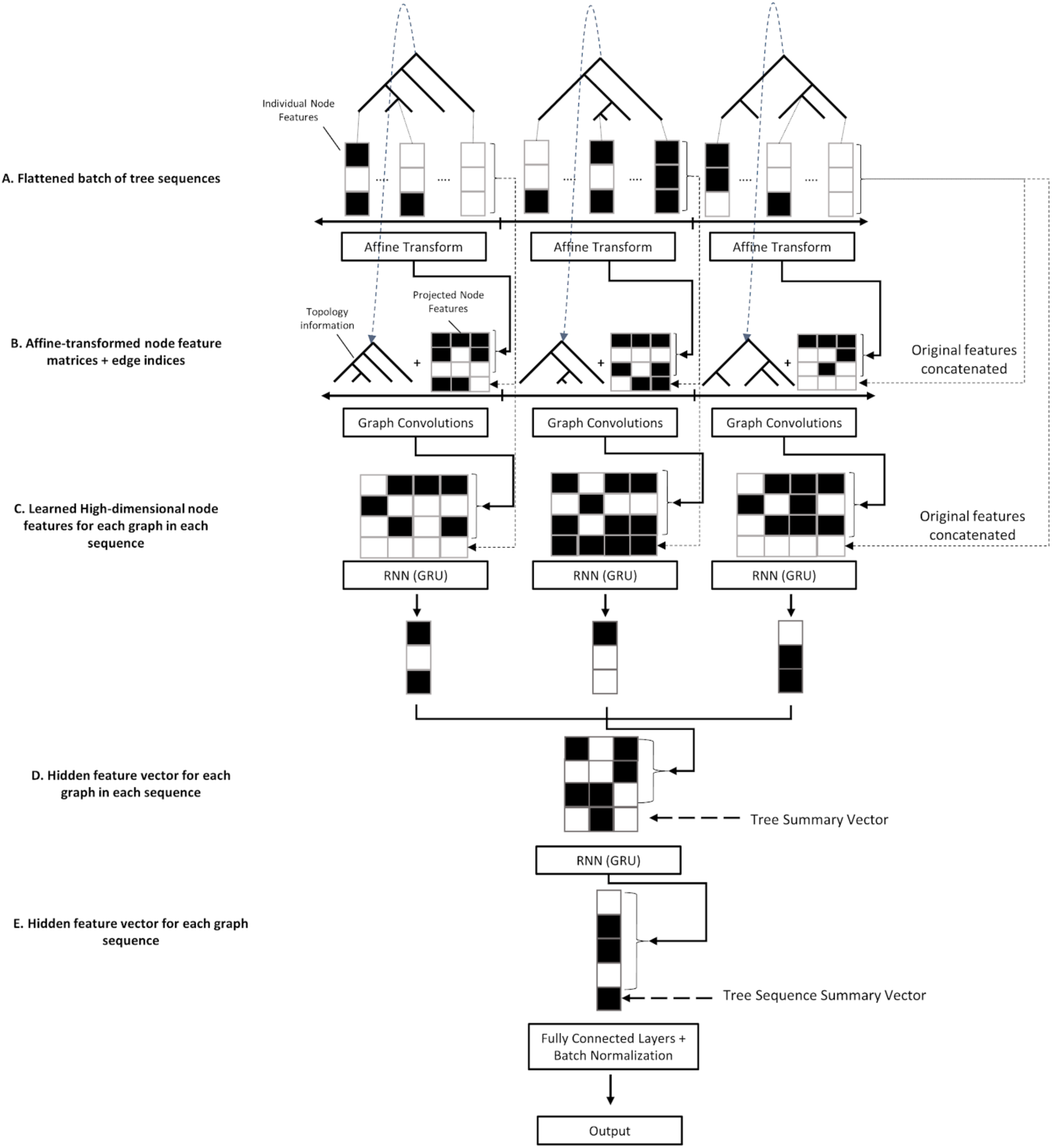
Overview of the GCN architecture. A) Each tree in the input tree sequence contains *n* nodes each summarized by *f* features, and is thus represented by an *n* × *f* node feature matrix per-tree, which is affine-transformed in the “node embedding” layer of the neural network (Table 5). B) For each tree, the affine-transformed node feature matrix is concatenated to a vector of edge indices specifying the tree topology, and also to the original feature matrix (i.e. a “skip connection”). This concatenated matrix is then input to multiple graph convolutional layers with skip connections. C) For each tree, the features from graph convolution layers are then fed as input to a GRU, resulting in a compressed, learned vector representation of the tree and its features. D) These compressed vector representations are concatenated to a “tree summary vector” that encodes summaries of the size and shape of the tree, as well as the size and location of the chromosomal segment corresponding to the tree. The resulting vectors for all trees are then concatenated together into one matrix corresponding to the entire tree sequence, and passed through another GRU layer that produces a flattened vector representation of the entire tree sequence. E) This vector is then concatenated to a “tree-sequence summary vector” containing information about the distribution of values in the original tree summary vectors and the number of trees in the original sequence (which may be less than the padded length of this vector). This vector is then passed through multiple fully connected layers culminating in a final output layer tailored to the target task. Throughout the diagram, solid lines with arrows show the flow of information through layers that transform the input, while dashed lines with arrows show the flow of information that has been unaltered by any preceding layers in the network.

### GCNs can leverage tree-sequence data to accurately estimate recombination rates

Estimating historical recombination rates is a deeply important task to population genetics as the recombination rate modulates key evolutionary processes and also affects our ability to make inferences about these processes. For example, when considering the effects of selection, the recombination rate between loci influences the degree of selective interference between them (Hill and Robertson 1966). Moreover, in the case of positive selection, the signatures of selection are much more difficult to detect in the presence of high rates of recombination due to the rapid exchange of alleles, which dampens the hitchhiking effect by preserving more genetic diversity in the vicinity of the selected site (Shriner et al. 2003). Because the recombination rate has a direct influence on the correlation between alleles in a sample of genomes, population genomic data can in principle be used to get an accurate estimate of the recombination rate landscape across the genome (Hudson 2001; Auton and McVean 2007; Chan et al. 2012; Gao et al. 2016; Adrion et al. 2020).

To assess the utility of tree-sequence based recombination rate inference, we trained our GCN for this task and compared its performance to the CNN previously shown by (Flagel et al. 2019) to accurately estimate recombination rates. Specifically, we simulated data using the same parameter values as in (Flagel et al. 2019) to generate samples of phased chromosomes which we used to estimate the crossover rate per base pair per meiosis by either passing these data directly into a CNN or through a tree-sequence inference procedure before passing the resulting trees into the GCN. In the context of tree sequences, recombination plays a very direct role in the structure of the data: most recombination events will alter the tree topology, and thus the number of trees in the tree sequence minus 1 is equal to the number of historical recombination events that could possibly be observed. However, because *inferred* tree sequences will contain errors, it is not clear whether we should expect methods that examine information from inferred tree sequences to estimate recombination to be more accurate than methods that directly examine the haplotype data. Indeed, it has been shown that ARG inference methods provide biased estimates of the number of recombination events (Deng et al. 2021). As shown in Figure 2, our GCN using *Relate*-inferred tree sequences were highly accurate, as were those produced by the CNN (Table 1) that takes the population genetic alignment as input. This provides support for the notion that a machine learning method trained on inferred tree sequences can learn to account for biases/errors present in these inferences—in this case a bias in the number of topology changes along the sequence. Thus, the presence of error in inferred tree sequences does not preclude the accurate inference of recombination rates from such data.

**Figure 2.**
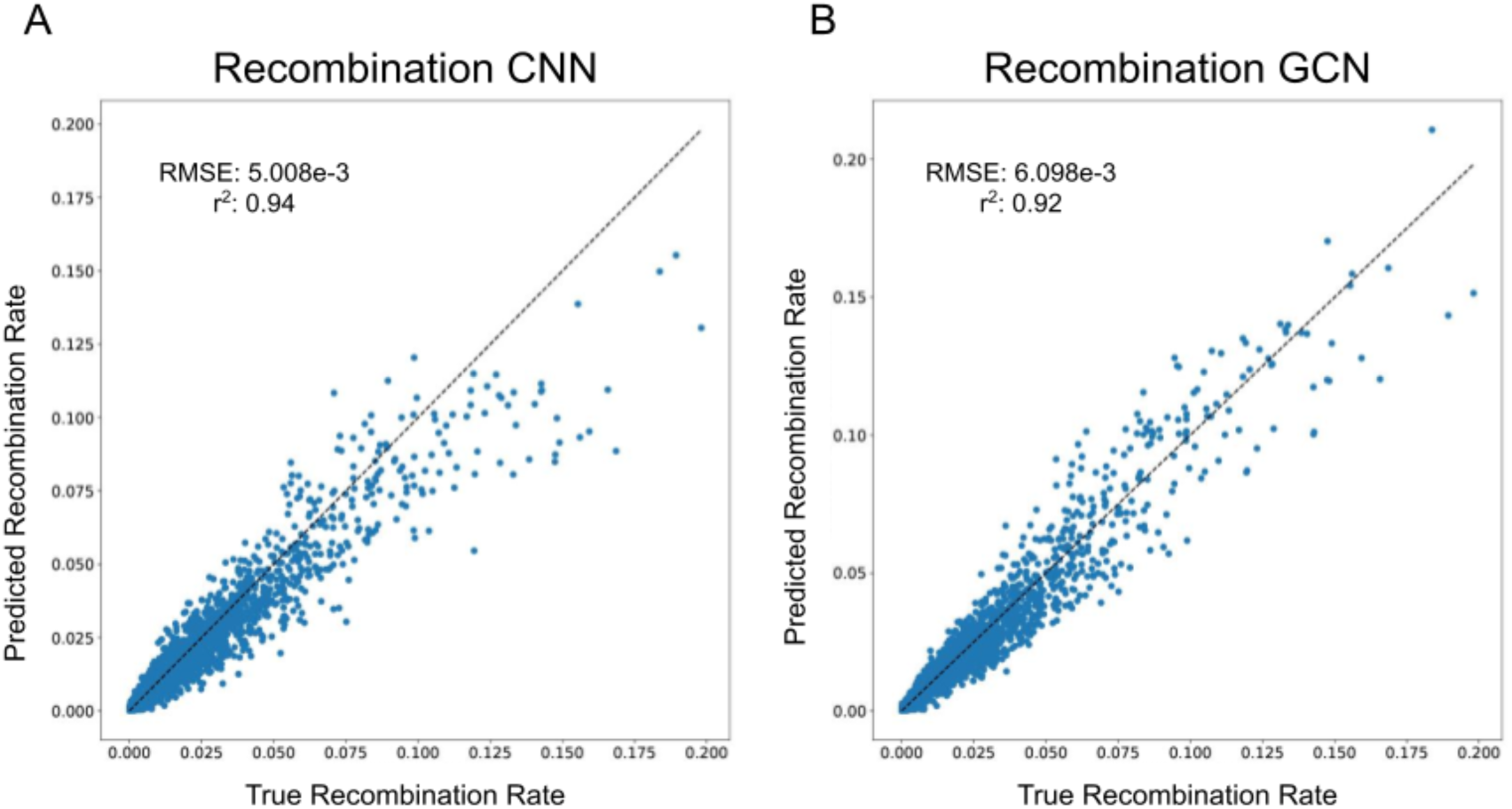
Predicted versus true recombination rates for the (A) CNN and (B) GCN architectures. Predictions are plotted against a reference diagonal dashed line, and each panel shows the root mean squared error (RMSE) and coefficient of determination (*R*^2^) for the given method.

**Table 1:** Loss and accuracy metrics for the CNN and GCN models across all tasks. For a given task, accuracy for the GCN and CNN were computed on the same test set. RMSE: root mean squared error; ROC AUC: area under the receiver operator correspondence curve; PR AUC: area under the precision-recall curve (i.e. average precision).

| Task | Metric | CNN | GCN |
| --- | --- | --- | --- |
| Recombination Rate Estimation | RMSE | **5.01E-03** | 6.10E-03 |
| Demography - $N_0$ | RMSE | 0.483 | **0.437** |
| Demography - $T_1$ | RMSE | 0.829 | **0.762** |
| Demography - $N_1$ | RMSE | 0.605 | **0.579** |
| Demography - $T_2$ | RMSE | 0.642 | **0.599** |
| Demography - $N_2$ | RMSE | **0.501** | **0.501** |
| Selection - Sweep vs Neutrality | ROC AUC | **0.91** | 0.825 |
| Selection - Sweep vs Neutrality | PR AUC | 0.706 | **0.857** |
| Selection - Hard vs Soft | ROC AUC | 0.856 | **0.86** |
| Selection - Hard vs Soft | PR AUC | 0.843 | **0.85** |
| Introgression - Introgression vs Neutrality | ROC AUC | 0.941 | **0.945** |
| Introgression - Introgression vs Neutrality | PR AUC | 0.975 | **0.977** |
| Introgression - Directionality | ROC AUC | 0.971 | **0.975** |
| Introgression - Directionality | PR AUC | 0.966 | **0.973** |

### GCNs dramatically outperform CNNs in detecting selective sweeps

The ability to detect loci affected by positive selection is a longstanding problem in population genetics (Stephan 2019). Selective sweeps occur when positive selection affecting an allele causes it to rapidly increase in frequency in a population, creating a valley of diversity that recovers at increasing genetic distances from the selected site (Smith and Haigh 1974; Kaplan et al. 1989). Sweeps also produce an excess of high-frequency derived alleles (Fay and Wu 2000), a deficit of haplotypic diversity (Hudson et al. 1994; Fay and Wu 2000; Sabeti et al. 2002), and elevated linkage disequilibrium (Kelly 1997; Kim and Nielsen 2004) near the target of selection. Recently, there has been great interest in characterizing and detecting the signatures of “soft sweeps” (Hermisson and Pennings 2017) which may be common in large, rapidly evolving populations (Hermisson and Pennings 2005; Karasov et al. 2010; Garud et al. 2015), and which produce somewhat different signatures than classic “hard” sweeps (Przeworski et al. 2005; Berg and Coop 2015). Recently, a number of machine learning methods have been designed to detect the multidimensional signatures of both of these types of sweeps (Pybus et al. 2015; Schrider and Kern 2016; Mughal and DeGiorgio 2019; Caldas et al. 2022; Hejase et al. 2022; Lauterbur et al. 2022; Whitehouse and Schrider 2022; Whitehouse and Schrider 2023).

However, it should be noted that the presence of all of these signatures of selection is a side effect of the profound effect that positive selection has on genealogies in the genomic vicinity of the selected locus. The shapes and sizes of trees are affected by selection in that the genealogies of loci near the selected site have a very short branches, especially internal branches, while genealogies at intermediate distances from the sweeping allele may have a subset of long branches as a consequence of recombination during the sweep (Fay and Wu 2000; Przeworski et al. 2005; Stephan 2019). Moreover, because of the reduced time period for historical recombination events to occur during the sweep, trees near the selected locus will correspond to a larger stretch of sequence than those from more distant loci that have experienced more recombination (Sabeti et al. 2002; Przeworski et al. 2005; Ferrer-Admetlla et al. 2014; DeGiorgio and Szpiech 2022). Thus, tree sequences may provide a great deal of information about recent positive selection.

To assess the potential of GCNs for detecting sweeps, we used the same simulated benchmarking scenario as (Flagel et al. 2019), and trained our networks to classify regions as undergoing a recent hard selective sweep (from a *de novo* mutation), being linked to a recent hard sweep, undergoing a soft sweep (from standing variation), being linked to a soft sweep, or evolving neutrally. Specifically, we simulated data under the JPT model from (Schrider and Kern 2017) to generate training, testing, and validation sets as described in the Methods. We find that the GCN architecture outperforms the CNN for all classes except neutrality, where their performance is equal (Figure 3). Indeed, the GCN obtains an average increase of 6.7% of test samples assigned to the proper class (the confusion matrix diagonal) relative to the CNN (65.5% accuracy for the GCN vs 58.8% for the CNN). Perhaps more strikingly, the GCN exhibits a sizable increase relative to the CNN in the area under the receiver operating characteristic (ROC) curve (AUROC) and in average precision (AP, equal to the area under the precision-recall curve) for distinguishing sweeps from neutrality (difference in AUROC of 0.085 and difference in AP of 0.151; Figure 4A, B). The increase in accuracy is less notable for distinguishing between hard and soft sweeps, with an AUROC increase of 0.004 and AP increase of 0.007 for the GCN compared to the CNN (Figure 4C, D).

**Figure 3.**
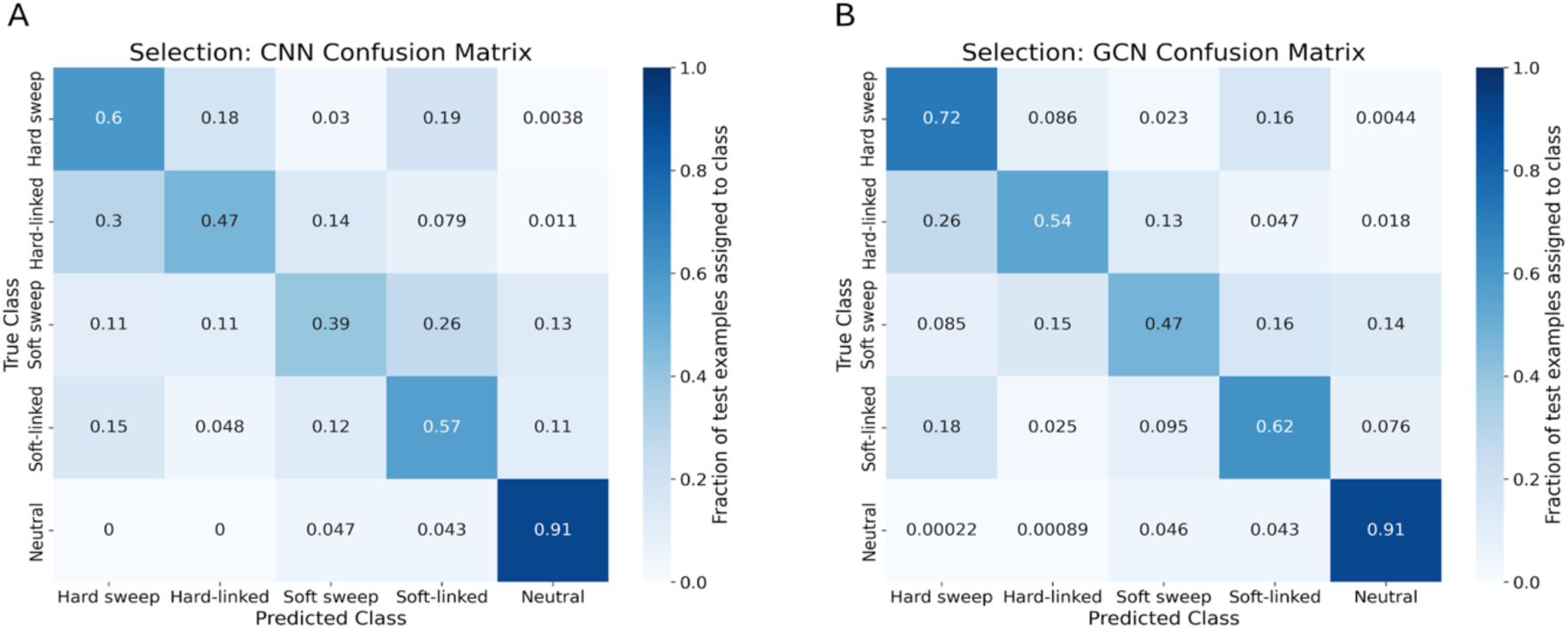
Confusion matrices showing performance of the CNN architecture and GCN architecture on the sweep detection task. A) Confusion matrix summarizing the CNN’s performance on a held-out test set. For this problem, there are five classes specifying the type of sweep (hard versus soft), whether a focal window is impacted by direct selection within the window or linked selection outside of the window (sweep vs. linked), or neutrally evolving. Each entry in the matrix shows the fraction of simulated examples under a given class (the True Class, specified by the row of the matrix) that were assigned to a given class by the network (the Predicted Class, specified by the column of the matrix). Diagonal entries thus represent correct classifications, and errors are represented in the off-diagonal entries, and each row sums to 1. B) Confusion matrix summarizing the GCN’s performance on the same test set.

**Figure 4.**
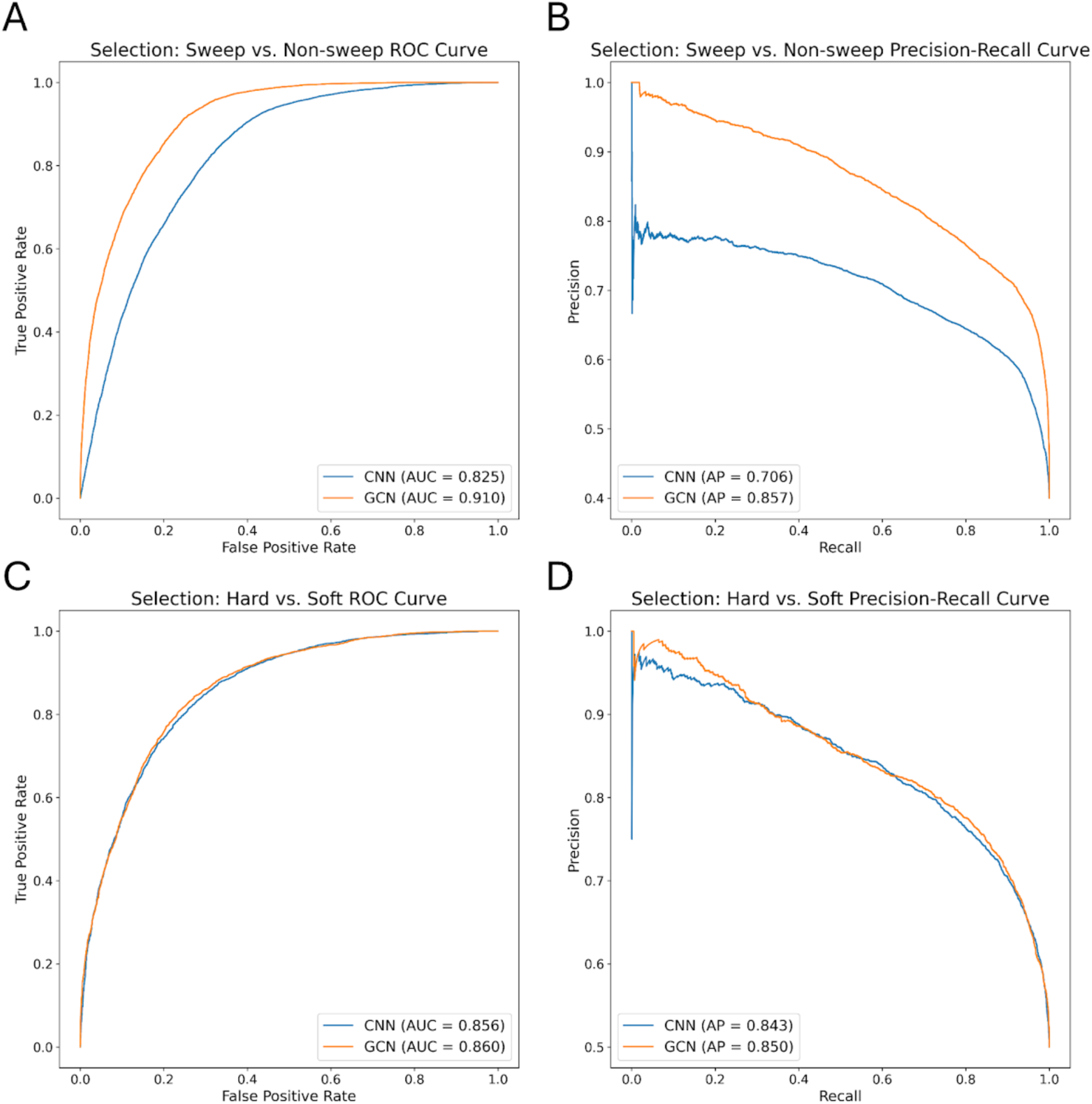
Receiver operating characteristic (ROC) and precision-recall (PR) curves summarizing performance of the CNN (blue) and GCN (orange) on the sweep-detection task. (A) ROC curves showing how well each method distinguishes between selective sweeps (whether hard or soft) and unselected regions (whether sweep-linked or neutral). The curves show the true and false positive rates for the CNN (blue) and GCN (orange) at varying classification thresholds. The score used to classify a window as a sweep vs. unselected at a given threshold was the sum of the neural network’s class membership probabilities for the “hard sweep” and “soft sweep” classes to yield a combined “sweep” probability. The area under the ROC curve (AUC) for each network is shown in the inset. B) Precision-recall curves showing the two network’s performance on the same “sweep vs. unselected” task, here showing the precision (i.e. the positive predictive value, or fraction of regions classified as sweeps that truly were sweeps) and recall (the true positive rate) at varying classification thresholds. The average precision (AP, which is equal to the area under the PR curve) is shown in the inset. C) ROC curves for distinguishing between hard and soft sweeps, with hard sweeps treated as positives and soft sweeps as negatives for this binary classification task. D) Precision-recall curves for the “hard vs. soft” task.

### GCNs perform similarly to CNNs on the task of detecting introgression

A variety of approaches have been developed to identify the signatures of introgression between pairs of closely related species (Hudson et al. 1992; Huson et al. 2005; Rosenzweig et al. 2016), including several machine learning methods (Schrider et al. 2018; Schrider et al. 2018; Gower et al. 2021; Ray et al. 2023). Previously, Flagel et al. (2019) showed that a CNN substantially outperforms FILET, a supervised machine learning method that detects introgression between closely related species with impressive accuracy (Schrider et al. 2018). It has also been shown that signatures of introgression can be extracted from inferred ARGs (Deng et al. 2024), suggesting that our GCN approach may work well on this task. We therefore compared the performance of the tree-sequence based GCN to our CNN on simulated data modeling gene flow between *Drosophila simulans* and *Drosophila sechellia* (see Methods for details). We find that both networks perform well (Figures 5,6), with the GCN slightly outperforming the CNN (86.2% accuracy for the GCN versus 85.1% for the CNN). The overall error profile is very similar for the two architectures (Figure 5) indicating that the GCN’s improvement over the CNN is not due to any specific class, but rather due to slight accuracy increases (1–2%) across all three classes.

**Figure 5.**
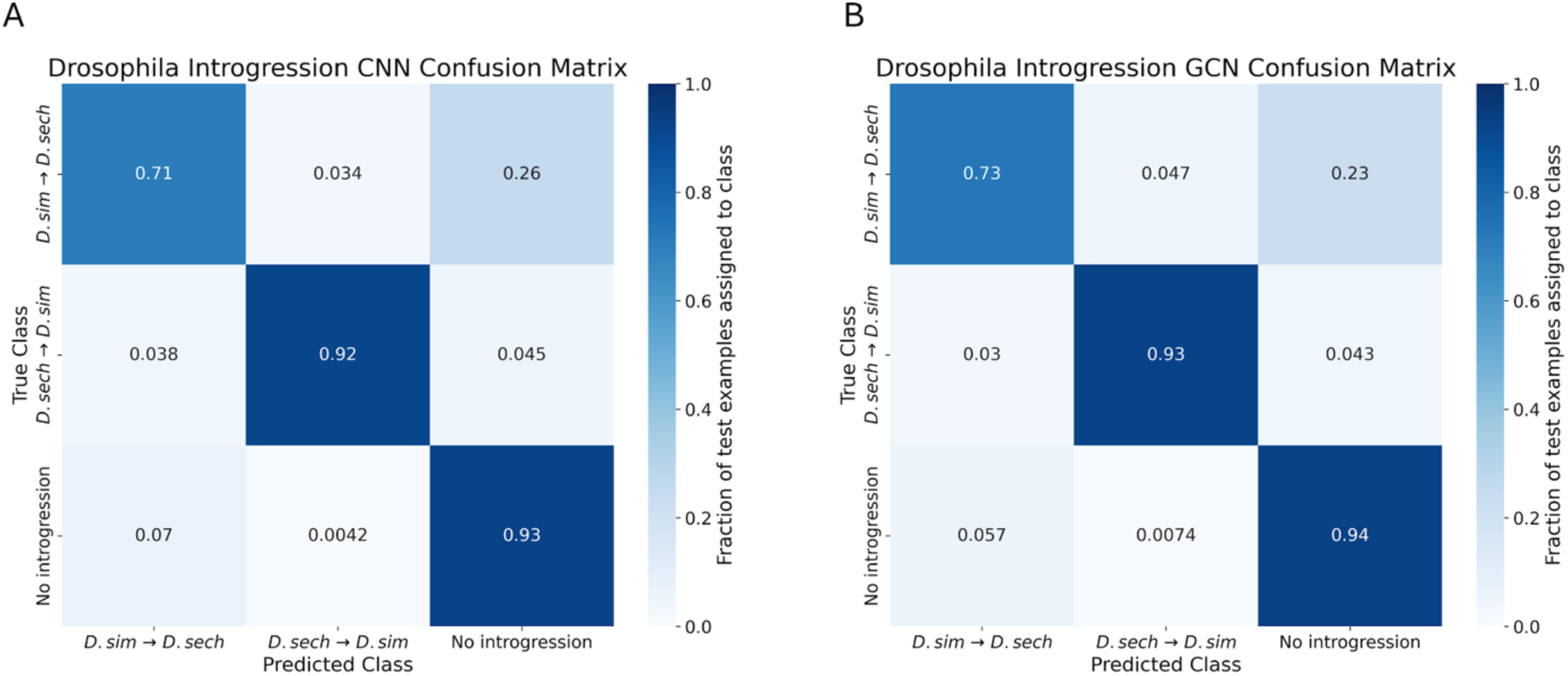
Confusion matrices showing the neural network’s accuracies on the introgression inference task. Confusion matrices are shown in the same manner as for Figure 3, but here the three classes are introgression from *Drosophila simulans* to *D. sechellia*, introgression from *D. sechellia* to *D. simulans*, and no introgression. A) Confusion matrix for the CNN. B) Confusion matrix for the GCN.

**Figure 6.**
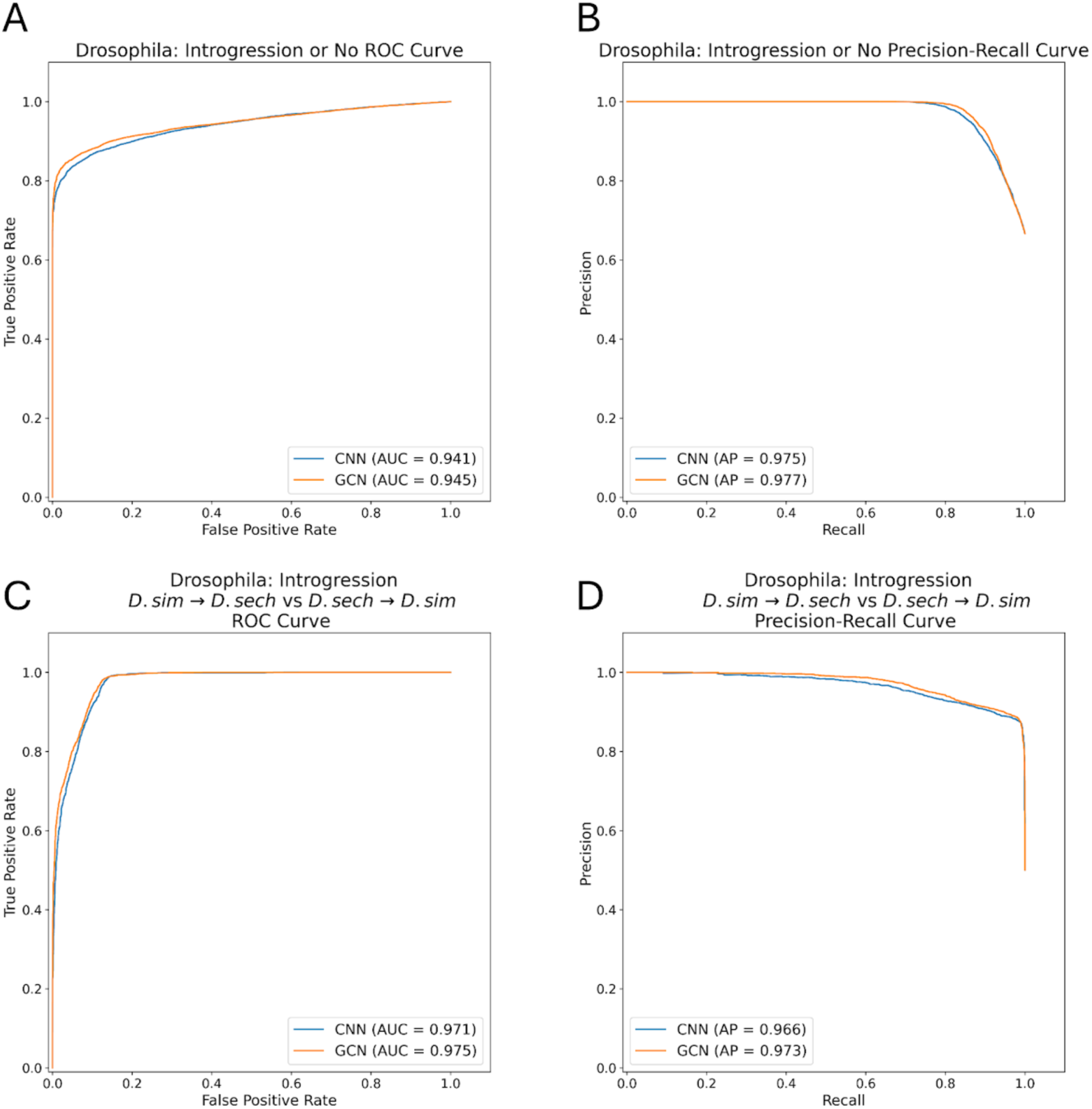
Receiver operating characteristic (ROC) and precision-recall (PR) curves for the introgression detection task. A) ROC curves summarizing the performance of the CNN (blue) and GCN (orange) on the task of distinguishing between windows with introgression (in either direction) and without introgression. The area under the ROC curve (AUC) for each network is shown in the inset. B) Precision-recall curves for the task of distinguishing between windows with and without introgression. The average precision (AP, which is equal to the area under the PR curve) is shown in the inset. C) ROC curves for the task of distinguishing between introgression from *D. simulans* to *D. sechellia* and introgression from *D. sechellia* to *D. simulans*. D) Precision-recall curves for the task of inferring the direction of introgression.

The GCN has an AUROC increase of 0.004 and AP increase of 0.002 when measuring accuracy distinguishing between genomic windows with and without introgression, and an increase in AUROC of 0.004 and AP of 0.007 when inferring the direction of introgression (Figure 6).

### GCNs show improvement compared to CNNs on a demographic inference task

Perhaps one of the most widely studied tasks in population genetics is the inference of demographic models and their associated parameters from. Many statistical approaches have been developed to allow researchers to estimate a population’s history of size changes, splitting and merging events, and expansions and contractions (Gutenkunst et al. 2009; Gutenkunst et al. 2009; Gutenkunst et al. 2010; Li and Durbin 2011; Browning and Browning 2015; Terhorst et al. 2017; Kamm et al. 2020; Santiago et al. 2020; Excoffier et al. 2021) These include several deep learning methods that have been designed in recent years (Sanchez et al. 2021; Wang et al.

2021), one of which applies graph convolutions to the tree sequence for inference (Korfmann, Sellinger, et al. 2023). To compare the performance of our GCN to the CNN on a demographic inference task, we reexamined the 3-epoch model presented in (Flagel et al. 2019). We simulated data under this model and assessed each network’s accuracy in estimating its 5 parameters: the population sizes in each of the 3 epochs, and the 2 transition times between these epochs.

We found that the GCN architecture outperforms the CNN on all estimations except the ancestral population size (*N*2) where the two methods perform equally as measured by both root mean squared error (RMSE) and the coefficient of determination (*R*^2^) on test data (Figure 7, Table 1). With respect to population size inference, the difference in accuracy between the GCN and CNN was most pronounced for the present-day population size (*N*0; improvement in RMSE and *R*^2^ of 0.046 and 0.048, respectively, for the GCN over the CNN), with the middle epoch’s population size inference (*N*1) showing smaller differences in accuracy (difference in RMSE and *R*^2^ of 0.026 and 0.045, respectively), and again no difference between the two methods for the ancestral size. For the inferred times of population size changes, we again observed greater difference in accuracy for the inferred time of the most recent event (*T*1; difference in RMSE and *R*^2^ of 0.067 and 0.069, respectively) than for the inferred time of the more ancient size change (*T*2; difference in RMSE and *R*^2^ of 0.043 and 0.047). This improvement in accuracy is somewhat modest (average RMSE decrease of 0.055 and average *R*^2^ decrease of 0.058 across all parameters), and we remind the reader that this test case involves the inference of demographic parameters from a single genomic region as a proof of concept rather than examining genome-wide data as is typically done in practice. Nonetheless, these results demonstrate the GCN’s ability to examine *Relate*-inferred tree sequences and extract information relevant to past demographic events at least as well as a CNN directly examining the population genetic alignment.

**Figure 7.**
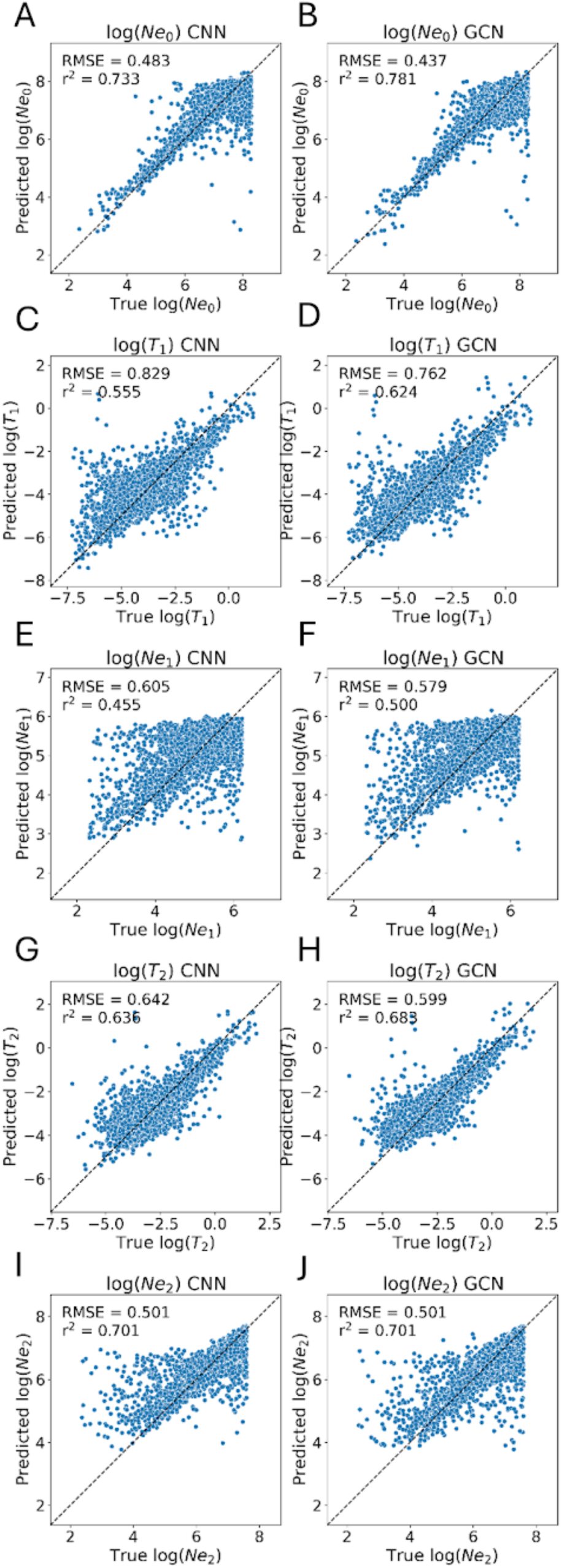
Performance of the CNN and GCN on the task of demographic parameter estimation on a held-out test set. The top row shows the predicted versus true values for the CNN (panel A) and GCN (panel B) for the present-day population size *N*0, and subsequent rows show the same for the time of the most recent population size change (*T*1, panels C and D), the second population size (*N*1, panels E and F), the time of the more ancient population size change (*T*2, panels G and H), and the ancestral population size (*N*2, panels C and D). Each panel also shows the root mean squared error (RMSE) and coefficient of determination (*R*^2^) for the given method on the given parameter. Values are log-scaled for visual clarity and plotted against a reference diagonal dashed line.

### Performance and computational efficiency

Perhaps the most enticing attribute of tree sequences is that they can provide extremely compact representations of population genomic data (Kelleher et al. 2019), leading to the possibility that they can be leveraged to reduce the computational burden of evolutionary inference pipelines.

We therefore compared the computational cost of our GCN and CNN on the problems examined above. With respect to preprocessing, our GCN’s pipeline has an advantage over our CNN pipeline in that tree sequences are relatively fast to infer compared to the sorting and matching steps that we perform on alignments before passing them into the CNN. For instance, computing *Relate* outputs for 1000 simulated replicates of length 10 kb from the introgression case took on average ∼ 5 minutes for a single CPU core vs. ∼ 21 minutes to seriate and match the same number replicates. In terms of memory efficiency, the size of a single replicate from the introgression data set after the tree-sequence inference step by *Relate* was 3.25 times the size of the original simulations (Tables 2, 3). Similar trends are observed for the recombination and demography problems. For sweep detection, the GCN requires slightly less memory than the CNN, but this is a consequence of our downsampling of the tree sequence for the GCN (Methods), while the CNN examines the entire input alignment. The size of a replicate is relevant because it influences memory capacity and speed of training. We note that it is possible that the memory requirements for the tree sequences would change substantially were we to use a different inference method. In addition, we note that the demographic inference problem required a smaller chromosome size than is typically used for such tasks (i.e. genome-scale data) due to the difficulty of storing an entire chromosome’s worth of information on GPU memory.

**Table 2:** GPU training and approximate RAM usage for the GCN architecture. All training times are reported for a single NVIDIA A100 GPU. RAM estimates were obtained via *nvidia-smi* while training was running. For the GCN, batch size in terms of number of input trees varies stochastically due to the variable tree sequence length.

| Problem | Time | RAM (1 batch) | RAM (1 replicate) | Batch size | Total epochs |
| --- | --- | --- | --- | --- | --- |
| Recombination rate | 7.9 hrs | 4.9 Gb | 38.3 Mb | 16 | 92 |
| Introgression | 12.4 hrs | 8.3 Gb | 231 Mb | 36 | 43 |
| Sweeps | 24.6 hrs | 10.7 Gb | 535 Mb | 20 | 67 |
| Demography | 14.5 hrs | 8.4 Gb | 280 Mb | 30 | 100 |

**Table 3:**
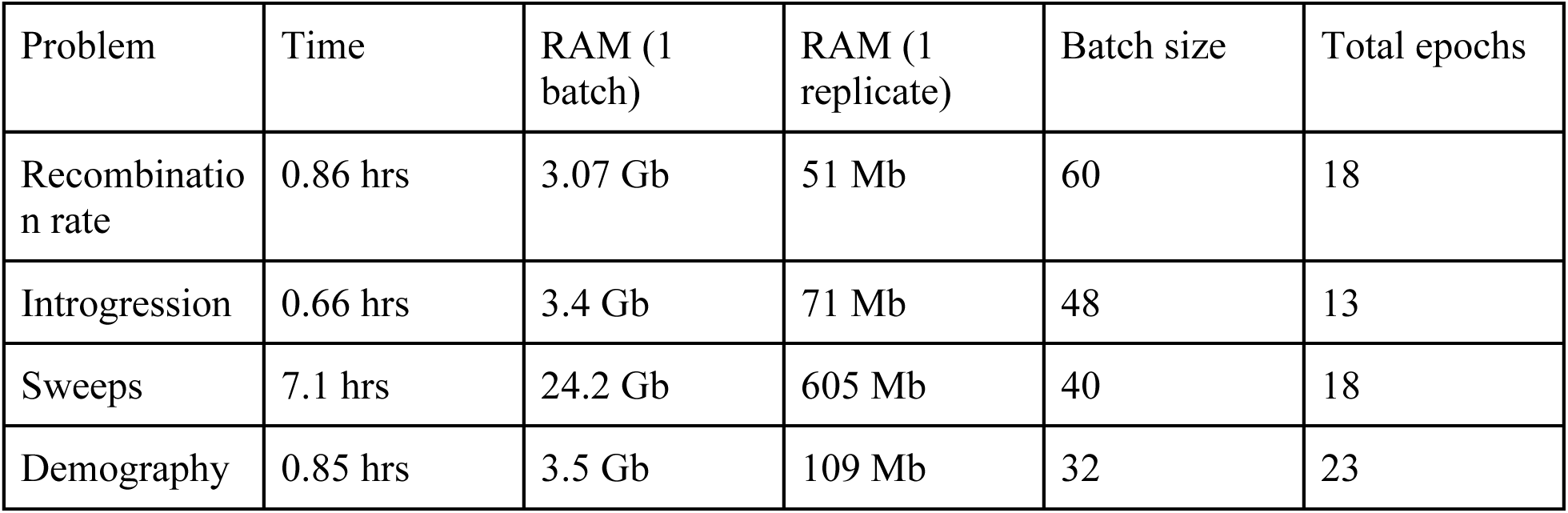
GPU training and approximate RAM usage for the CNN architecture. All training times are reported for a single NVIDIA A100 GPU.

| Problem | Time | RAM (1 batch) | RAM (1 replicate) | Batch size | Total epochs |
| --- | --- | --- | --- | --- | --- |
| Recombination rate | 0.86 hrs | 3.07 Gb | 51 Mb | 60 | 18 |
| Introgression | 0.66 hrs | 3.4 Gb | 71 Mb | 48 | 13 |
| Sweeps | 7.1 hrs | 24.2 Gb | 605 Mb | 40 | 18 |
| Demography | 0.85 hrs | 3.5 Gb | 109 Mb | 32 | 23 |

Nonetheless, because the same simulated data was used for both the GCN and the CNN, the performance comparison included here is a fair one.

Training time is also an important consideration when choosing deep learning architecture. We note that our CNN, due to the smaller input sizes and the greater efficiency of image convolutions over graph convolutions with current hardware and software, are faster to train and usually require less RAM (Tables 2, 3). This is in spite of the fact that our choice of CNN has almost 2 orders of magnitude more learnable parameters than our proposed GCN architecture (21,280,965 vs. 228,837 respectively). However, we find both approaches to be accomplishable on consumer-grade hardware, i.e. on a single GPU with ∼ 12 Gb of RAM.

We note that, in addition to training the network weights themselves, part of the computational cost of each method is to preprocess the training data. For the CNN, the preprocessing step is to sort the input genotype matrix, which has previously been shown to improve accuracy (e.g. Flagel *et al*. 2019; Ray *et al*. 2023). For the GCN, the tree inference step itself must be performed on the training data. We therefore compared the amount of time required to preprocess the input data for both the GCN and CNN architectures. We found that tree-sequence inference for the GCN was faster than input sorting for the CNN for each task (Table 4). We note that seriation is a computationally expensive process, sometimes a factor of 10 times more per chromosome than tree sequence inference (Table 4). However, we note that, for the CNN’s preprocessing step, one could potentially identify a more computationally efficient sorting algorithm than seriation that could still yield acceptable performance. Finally, we stress that these training steps incur a one-time cost, as once the network is trained it can be used to perform rapid inference on each instance on an entire data set (e.g. infer the recombination rate in each of thousands of sliding windows in the genome) with no additional training.

**Table 4:** Preprocessing computation time per method. Speed is in units of seconds per chromosome. Times were each computed using a single Intel Core i9-9900K CPU core clocked at 3.60GHz.

| Problem | Relate (s / chrom) | Sorting (s / chrom) | Sorting algorithm | Number of sampled individuals |
| --- | --- | --- | --- | --- |
| Introgression | 0.11217 | 1.00905 | Seriation + linear sum assignment | 34 |
| Selection | 1.0812 | 1.1217 | Seriation | 104 |
| Recombination | 0.04866 | 0.7921 | Seriation | 50 |
| Recombination (large) | 1.367 | 1.183 | Seriation | 100 |
| Demography | 0.1239 | 0.9647 | Seriation | 50 |

**Table 5:**
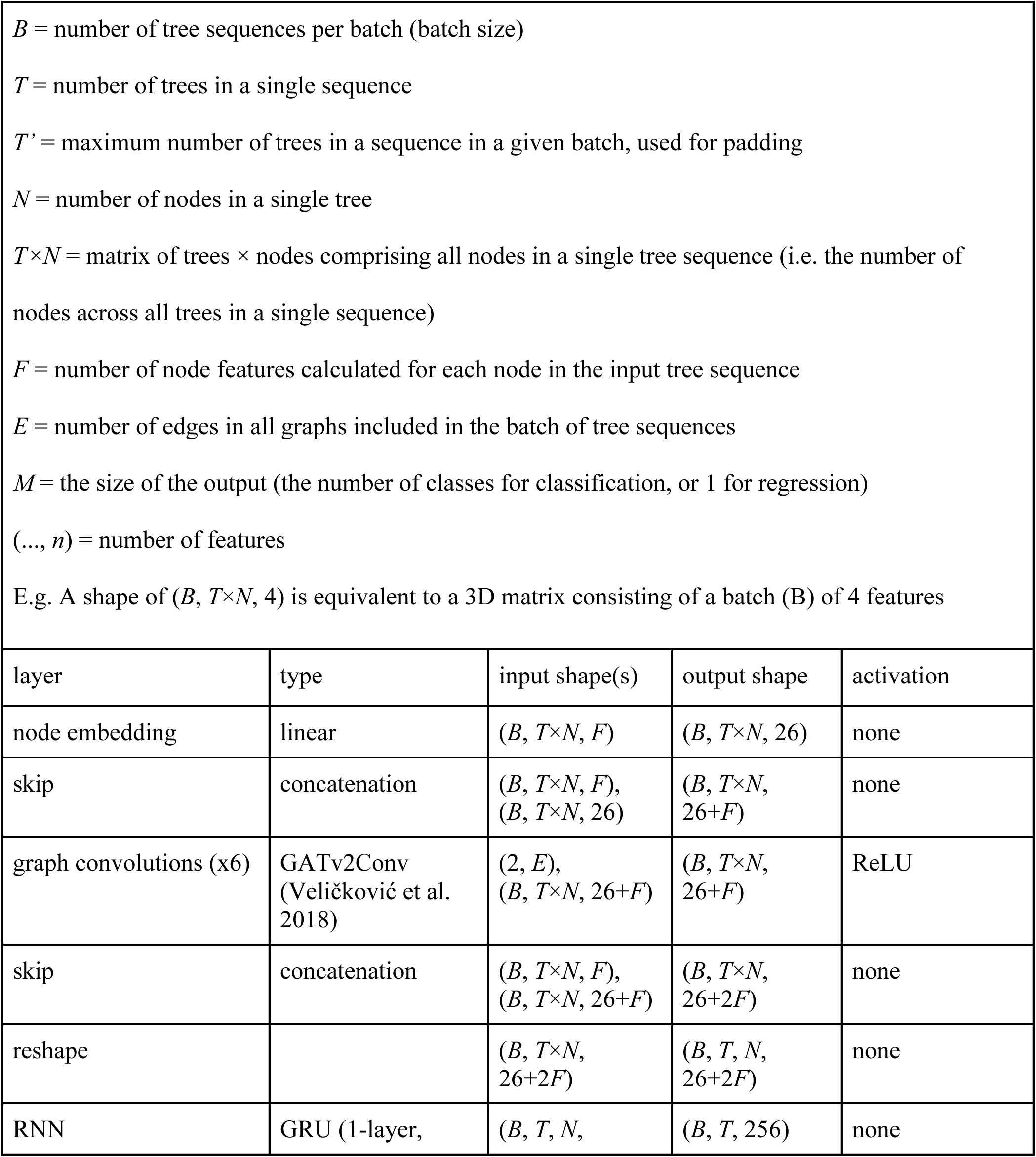
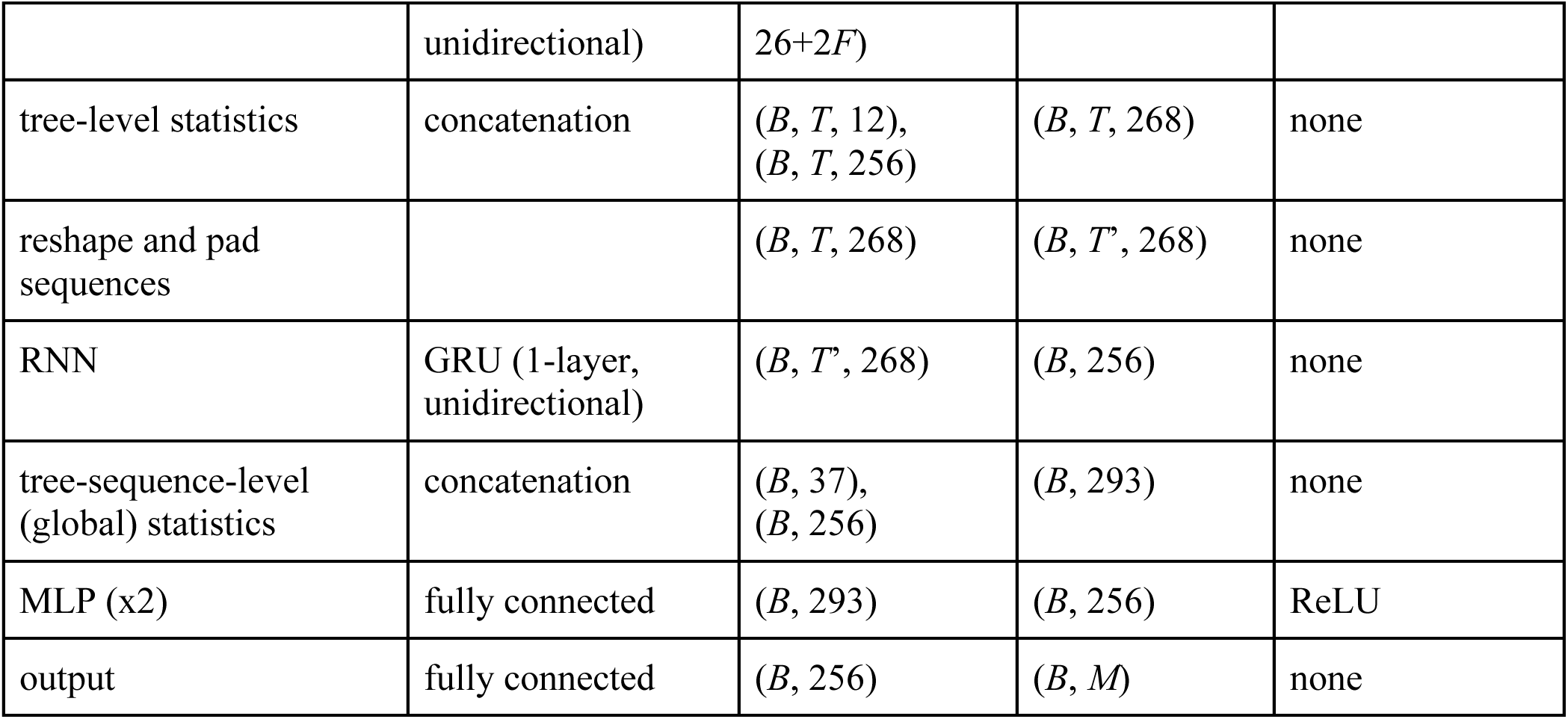
Graph neural network layers and their input/output shapes and activation functions.

| $B$ = number of tree sequences per batch (batch size)<br>$T$ = number of trees in a single sequence<br>$T'$ = maximum number of trees in a sequence in a given batch, used for padding<br>$N$ = number of nodes in a single tree<br>$T \times N$ = matrix of trees × nodes comprising all nodes in a single tree sequence (i.e. the number of nodes across all trees in a single sequence)<br>$F$ = number of node features calculated for each node in the input tree sequence<br>$E$ = number of edges in all graphs included in the batch of tree sequences<br>$M$ = the size of the output (the number of classes for classification, or 1 for regression)<br>$(..., n)$ = number of features<br>E.g. A shape of $(B, T \times N, 4)$ is equivalent to a 3D matrix consisting of a batch ($B$) of 4 features | | | | |
| --- | --- | --- | --- | --- |
| layer | type | input shape(s) | output shape | activation |
| node embedding | linear | $(B, T \times N, F)$ | $(B, T \times N, 26)$ | none |
| skip | concatenation | $(B, T \times N, F),$<br>$(B, T \times N, 26)$ | $(B, T \times N,$<br>$26+F)$ | none |
| graph convolutions (x6) | GATv2Conv<br>(Veličković et al.<br>2018) | $(2, E),$<br>$(B, T \times N, 26+F)$ | $(B, T \times N,$<br>$26+F)$ | ReLU |
| skip | concatenation | $(B, T \times N, F),$<br>$(B, T \times N, 26+F)$ | $(B, T \times N,$<br>$26+2F)$ | none |
| reshape | | $(B, T \times N,$<br>$26+2F)$ | $(B, T, N,$<br>$26+2F)$ | none |
| RNN | GRU (1-layer, | $(B, T, N,$ | $(B, T, 256)$ | none |
| | unidirectional) | $26+2F$ | | |
| tree-level statistics | concatenation | $(B, T, 12),$<br>$(B, T, 256)$ | $(B, T, 268)$ | none |
| reshape and pad sequences | | $(B, T, 268)$ | $(B, T', 268)$ | none |
| RNN | GRU (1-layer, unidirectional) | $(B, T', 268)$ | $(B, 256)$ | none |
| tree-sequence-level (global) statistics | concatenation | $(B, 37),$<br>$(B, 256)$ | $(B, 293)$ | none |
| MLP (x2) | fully connected | $(B, 293)$ | $(B, 256)$ | ReLU |
| output | fully connected | $(B, 256)$ | $(B, M)$ | none |

### Assessing sensitivity to misspecification

The robustness of population genetic inference models to misspecification of the assumed demographic model or its parameters is an important consideration as in practice both are inferred and never directly observed (Schrider and Kern 2016; Johri et al. 2022; Mo and Siepel 2023). To test how sensitive each method is to misspecification, we generated two new sets of simulations for inferring the presence of selection. In the first, we applied the networks trained on our original training set that includes population size changes to a test set with no history of population size change but otherwise identical parameters to see what decrease in performance, if any, is observed for each method. This misspecification of the demographic history resulted in a significant and similar decrease in accuracy for both methods with the GCN still outperforming the CNN. On this common evaluation set with no population size change (25,000 examples), the CNN achieved an accuracy of 36.3% (vs its original accuracy of 58.8%) while the GCN achieved an accuracy of 40.4% (vs its original accuracy 65.5%).

For the second set, we generated 50,000 simulations, in batches of 50, with each batch having different distributions of *θ* and *ρ* than the original training set. Specifically, for each batch we shifted the uniform range of *θ* values and the truncated exponential distribution *ρ* each relative to the original simulated distributions by separate multiplicative factors drawn uniformly from *U*(0.5, 1.5) (i.e. for *θ* both the lower and upper bounds of the original uniform range are multiplied by a value between 0.5 and 1.5, and for *ρ* both the mean and the upper bound of the original bounded exponential are multiplied by a different value between 0.5 and 1.5); this procedure results in a set that contains a mixture of a large number of parameter distributions that is much broader than that included in the original training set.

We first applied our original network to the new test set with a broader range of *θ* and *ρ*parameters. In this case, the CNN obtained better accuracy than the GCN (56.3% vs 46.1%), implying a great misspecification effect for the GCN. To confirm that this was the case, and to see whether either method’s performance is adequate when trained on a broader range of parameters that encompasses that of the test set, we re-trained each method on the combined superset of both the original and more variable training sets for a short number of epochs starting with the weights obtained prior. Here, the GCN outperformed the CNN 66.3% to 60.5%, confirming that the former suffers a greater drop in accuracy when applied to data outside of the training distribution. However, the GCN also takes as input summary statistics (e.g. the vector of all coalescence times in the tree) that are normalized according to values observed in the training set, and in practice one would be able to observe whether these values differ greatly between the training and test data and therefore re-normalize accordingly. We therefore asked how much of the GCN’s drop in performance could be remedied by keeping the neural network model the same but renormalizing the input: we used the same network weights (obtained on the misspecified training set) but normalized the input node times, tree summary vectors, and tree sequence summary vectors using the means and standard deviations calculated from the new training set. We found that performance increased to 52.5%, closer to that of the misspecified CNN but still well below that of the GCN trained on the broader range of parameters (Figure S1). Thus, correcting the summary statistic means and standard deviations appears to mitigate the misspecification effect somewhat but does not eliminate it entirely.

Finally, we note that for all of the simulated scenarios presented in this section, the original values for effective population size, and recombination and mutation rates that are passed to *Relate* are kept the same even when retraining on the data set that includes a larger more diverse distribution of these parameters. Thus, all of our results on this test data set also include the effect of a different sort of misspecification: differences between the parameter values given as input to *Relate* and the true values of these parameters. Overall, our GCN performs reasonably well in the presence of misspecification—although the magnitude of the reduction in accuracy caused by misspecification was higher for the GCN than for the CNN for both problems, the GCN’s accuracy remained substantially higher than that of the CNN in one of these cases despite the drop in accuracy due to misspecification. This result implies that the misspecification of tree sequence inference parameters may not have an especially drastic effect on accuracy, and for our GCN this effect can be mitigated by properly normalizing input summary statistics.

### Assessing the impact of chromosome and sample size

To test what impact chromosome length and the number of samples has on performance, we simulated additional training for the recombination regression problem with the same distribution of parameters as before but with twice the original sample size (100 individuals instead of 50) and 10 times the original chromosome length (200 kb instead of 20 kb). After training in the same way as described in the Methods, we again evaluated both the CNN and GCN on a common evaluation set. We found that both approaches accuracy when predicting *ρ* or the scaled recombination rate increases significantly with the GCN achieving an RMSE of 3.62×10^-3^ and the CNN again achieving marginally better accuracy at 3.45×10^-3^.

### The impact of neural network depth on performance

We sought to assess the impact of the depth of the GCN and CNN architectures on performance by altering the depth of these networks and then retraining them to infer recombination rates using the same data we had simulated prior. Specifically, we trained both the CNN and GCN with three different numbers of convolutional layers. In the case of the GCN, we trained with 6 (our choice for experiments in this work), 12, and 24 graph convolution layers in the network. For the CNN we used three variants of the ResNet architecture, ResNet34 (our choice), ResNet50, and ResNet101 (Figure S2). We find that these architectural choices did not result in sizable differences in accuracy, outside of the early training epochs. Thus, given that our 6-layer GCN and a ResNet34 CNN did not perform worse than deeper networks, our architectural design choices appear to be acceptable.

### The impact of removing GCN architecture components

We experimented with removing certain components in our chosen GCN architecture to assess their contribution to the GCN’s performance. This was done by re-training our GCN architecture to infer recombination rate using the same training set but with one or more parts of the network missing. First, we removed all graph convolutions leaving only two GRUs operating over the node and graph level features. Next, we experimented with retaining the graph convolution layers, but removing local features (the “tree summary vector” detailed in Methods), global features (or the “tree sequence summary vector”), or both sets of features from input to the network. Figure S3 shows the validation loss curves over training for each modified network architecture choice. We find that graph convolution or message passing over the inferred tree topologies is highly beneficial to the network’s ability to infer recombination rates—the removal of graph convolutions produced the largest increase in the minimum validation loss achieved during training. Without graph convolution, the topology is ultimately absent from the input to the network, but the network is still able to use the estimated times of ancestral nodes and mutations on the parent branches. If we instead retain the graph convolutions but remove either the local or global features we observe only a small decrease in accuracy, although training stability somewhat suffers. Finally, removing both global and local features (i.e. using features extracted by the graph convolution layers only) leads to very unstable training and a minimum loss that is somewhat higher than that obtained when including the global and local features, but still substantially lower than that achieved when using only these features and omitting the graph convolutions. Thus, while overall it appears that the graph convolution layers have the largest impact, each type of feature included in our network seems to improve performance.

### Using an alternative tree sequence inference method

We sought to observe whether using a different inference tool to obtain tree sequences from simulated data would affect the performance of our GCN. To do so, we applied *tsinfer* (Kelleher et al. 2019) to the training set for recombination rate estimation, created the GCN’s input from *tsinfer*’s output in the same manner as we did for *Relate* (Methods), and then trained our GCN architecture on these data. In contrast to *Relate, tsinfer* takes as input only the recombination rate, and does not use an estimate of effective population size or mutation rate.

After re-trainining our GCN using trees inferred by tsinfer, we applied the network to the same evaluation we used above to compare the GCN and CNN. We found that using *tsinfer* led to similar but slightly lower recombination rate estimation accuracy: we obtained an RMSE of 6.31×10^-3^ (vs. the *Relate*-trained GCN’s RMSE of 6.01×10^-3^) and an *R*^2^ value of 0.92 (the same value obtained by the *Relate*-trained GCN). While using *Relate* resulted in slightly better accuracy than *tsinfer* for this problem, others have found the opposite in the case of demographic inference (Fan *et al*. 2023). Thus, while our results demonstrate that our GCN approach can be readily and successfully extended to use alternative tree sequence/ARG inference methods, further would be necessary to characterize the interplay between inference method, estimation task, and network architecture with respect to inference accuracy.

## DISCUSSION

The size and quantity of population genetic datasets are increasingly large, making downstream analysis more computationally intensive. As researchers develop new, more compact data structures for population genetics, new methods must be developed to fully take advantage of these structures’ potential to tackle long standing problems in the field with greater computational efficiency and perhaps better accuracy as well. Tree sequences have become increasingly popular, both in the context of estimation from empirical data and in simulation (Kelleher et al. 2018; Haller et al. 2019; Ralph et al. 2020). However, there has yet to be a fundamental shift in population genetics inference approaches away from less efficient data structures (Korfmann, Gaggiotti, et al. 2023).

In principle, tree sequences have a key advantage over genotype information in that they can represent the complete evolutionary history of a sample of genome sequences, making them an attractive avenue for downstream evolutionary inference. However, an important caveat with using tree sequences is that in empirical data the true evolutionary history of a sample can never be known, and therefore tree sequences must be estimated from genotypes (Kelleher et al. 2019; Mahmoudi et al. 2022). As it stands, inference methods can produce highly inaccurate tree sequences when benchmarked on simulated data (Zhang et al. 2023), suggesting that using tree sequences as input for further inferences may be problematic. Moreover, the paradigm of inferring a population sample’s tree sequence to use as input for a downstream inference task does on the face of it appear to be unnecessarily complicated, and appears to violate the following imperative from Vapnik: “When solving a problem of interest, do not solve a more general problem as an intermediate step” (Vapnik 2006). For example, if we wish to infer whether a given locus has experienced a selective sweep, it may not be desirable to first infer the full evolutionary history of every segment of DNA along a large stretch of chromosome surrounding this locus, as this would clearly be a more challenging task than directly searching for evidence of recent positive selection.

Why then, might tree sequence-based inference still be generally highly effective, as our results seem to suggest? First, it seems probable that incorrect tree sequences contain information that, although incorrectly describing the true evolutionary history of a sample, can be useful for population genetic inference. In addition, tree sequences allow for a more concise representation of population genetic data than a genotype matrix. Moreover, tree sequence inference may serve as a form of de-noising by aggregating data across sites in a region in order to extract a meaningful pattern that may not be easily observable directly from sequence data (especially in the presence of sequencing error), even if that pattern does not describe the true evolutionary history. In any case, supervised machine learning methods combined with tree sequences can be especially powerful because one can train the machine learning model on tree sequences that are inferred from simulated data (as we have done here), rather than the true tree sequences, so that the training tree sequences may exhibit a similar error profile as tree sequences inferred from real data. In this framework, it does not matter whether the tree sequences are correct, but only that they contain information useful for the task at hand. Moreover, the impact of any remaining fundamental differences between tree sequences from simulated vs. real data could potentially be mitigated via the machine learning technique of domain adaptation (Ganin and Lempitsky 2015), wherein a network can be trained to learn to ignore systematic differences between simulated and real data before downstream prediction (Mo and Siepel 2023); as an aside, domain adaptation could also help mitigate the effects of demographic misspecification that we observed. Thus, in spite of the somewhat unusual workflow, relying on inferred tree sequences as input to supervised machine learning methods may in fact prove to be an especially powerful and robust strategy.

It is important to note that machine learning may not always be essential for obtaining accurate estimates from inferred genealogies. Indeed, recent studies have shown that while performing statistical inference using inferred trees substantially increases the error rate of downstream inferences (e.g. a ∼3- to ∼6-fold increase in the error of demographic inferences by *gLike*; (Fan et al. 2023)), the resulting inferences are still quite accurate in comparison to competing methods (Stern et al. 2019; Fan et al. 2023; Link et al. 2023). On a related note, although we feel that our comparison of the GCN’s performance to that of the CNN was sufficient to demonstrate the general utility of population genetic inference based on estimated genealogies, we note that we have not compared our GCN to other ARG or tree sequence-based methods. We therefore cannot make any conclusions regarding the relative effectiveness of deep learning methods versus statistical approaches that directly examine inferred genealogies.

However, we again stress that the accurate recombination rate estimates from our GCN, in spite of biases in the number of tree topology changes in the input, suggests that an SML approach could potentially add important value in one key respect: the ability to flexibly account for any biases introduced by the genealogy inference software by training on such biased genealogies.

Our goal was to examine the effectiveness of using inferred population genetic tree sequences as input for downstream inference methods, and to this end we compared our GCN to a population genetix alignment-based CNN; the latter was previously shown to be a powerful general-purpose approach for making population genetic inferences directly from alignment data (Flagel et al. 2019). Although we have provided an attempt at comparing the effectiveness of the GCN and CNN on multiple population genetics inference tasks, this set of problems was of course far from comprehensive. Moreover, we examined only one architecture for each input type, in contrast to the rapidly expanding set of supervised machine learning methods tailored to specific population genetic inference tasks: in recent years numerous SML methods have been developed to detect natural selection (Kern and Schrider 2018; Hejase et al. 2022; Whitehouse and Schrider 2023; Riley et al. 2024), identify loci affected by introgression (Schrider et al. 2018; Gower et al. 2021; Ray et al. 2023), estimate recombination landscapes (Adrion et al. 2020) and detect hotspots (Chan, Perrone, Spence, et al. 2018), and infer organisms’ dispersal rates (Smith et al. 2023; Smith and Kern 2023), among others (reviewed in Schrider and Kern 2018, Korfmann *et al*. 2023a). Thus, we make no claims about the methods being tested here achieving the best performance on any particular problem. Nonetheless, the encouraging benchmarking results from our GCN demonstrate the general effectiveness of tree sequence- based inference, and suggest that machine learning methods using tree sequences as input will likely prove to be a powerful addition to the ever-growing suite of tools and problems described above. Indeed, the success of our GCN on several very different tasks suggests that graphical neural networks may be adept at constructing lower dimensional embeddings of genealogical representations of population genomic data. An exciting extension of our work would thus be to train a general-purpose embedding network that could be used to extract features for training task-specific SML methods without the need for the relatively computationally intensive training performed here.

One important limitation of our work is that, although our networks were able to be trained and executed on consumer-grade hardware, we have not made exhaustive attempts to improve computational efficiency. Indeed, we find that our CNN is more efficient than the GCN in terms of both memory usage and training time, with the current advantage of the GCN being its faster preprocessing time. However, the latter is a consequence of our choice to sort our alignments in order to improve performance, a computationally expensive process (Ray et al. 2023) and it is important to note that the cost of this step will depend on the sorting algorithm used and the size and number of alignments to be sorted. Moreover, at least some problems, exchangeable neural networks that do not require any sorting may be used (Chan, Perrone, Spence, et al. 2018). However, we note that the advantage in compactness afforded by tree sequences is not observed at small sample sizes (Kelleher et al. 2019), and the tasks examined here all used small to modest sample sizes. Thus, our conclusions about the relative efficiency of GCNs vs. CNNs may not hold for tasks requiring much larger samples. Further investigations into optimizations of both architecture and processing algorithms may increase the favorability of the GCN in all aspects. future efforts experimenting with more compact tree representations (Kelleher et al. 2018; Haller et al. 2019; Kelleher et al. 2019; Ralph et al. 2020; Mahmoudi et al. 2022) and different neural network architectures may yield improvements in this area. Another limitation of our study is that we primarily used a GCN trained on *Relate*-inferred genealogies, and while our accuracy did not change substantially when estimating recombination rates using input data inferred by *tsinfer* (Kelleher et al. 2019), it is possible that for some tasks the accuracy of downstream inference may be affected by the method used to infer the input trees. Lastly, while we chose to focus on the relative performance of our GCN and CNN to assess the general effectiveness of tree sequence-based inference, another class of methods that we did not examine are those that make statistical inferences directly on genealogies, such as CLUES, which estimates selection coefficients from ARGs (Stern et al. 2019), and gLike, which can calculate the likelihood of a demographic model given an observed genealogical tree (Fan et al. 2023).

Thus, we cannot make any claims about the performance of these methods relative to the neural networks considered here.

Finally, we note that we have only explored a relatively small number of population genetic inference tasks that are of particularly broad interest to researchers. We therefore encourage exploration of tree sequence-based deep learning inference as a general framework for population genetic inference. Such work could also include experimentation with additional deep learning strategies for graph-based population genetic inferences that could be explored, such as neural networks that can take advantage methods capable of inferring the full ARG rather than a sequence of marginal trees (e.g. Rasmussen *et al*. 2014; Terhorst *et al*. 2017; Mahmoudi *et al*. 2022), and process the inferred ARG as a single graph rather than a sequence thereof. This and other approaches could potentially yield greater gains than our GCN which extracts information from each tree in the ARG before combining this information with a recurrent neural network. Our hope is that additional tree sequence-based deep learning research will not only lead to improved accuracy on problems but also, as alignment-based deep learning applications have done, empower researchers to tackle new problems (Battey et al. 2021; Yelmen et al. 2021; Booker et al. 2023; Korfmann, Gaggiotti, et al. 2023; Smith et al. 2023; Whitehouse and Schrider 2023).

## METHODS

### Tree sequence inference

For each problem, simulations were performed as described in the sections below, producing *ms*- style output. The simulation output was then formatted for input to *Relate* (Speidel et al. 2019), a recently developed method that, when given a sample of genomes along with estimate of the mutation rate, recombination rate, and effective population size, infers the sample’s genome-wide genealogy in the form of a sequence of rooted binary trees. *Relate* outputs this sequence of trees in order along the chromosome, with each tree spanning a variable length of sequence bounded by the breakpoints of inferred historical recombination events that altered the tree topology. This was done for each simulation replicate for each problem described below, using the mutation, recombination rate and effective population size parameters listed in Table S1.

These parameters (mutation and recombination rate, and effective population size) are inputs to *Relate,* and thus an assumed value or mean value for these parameters is needed prior to inference. In each case in this paper where a distribution of values for these parameters is simulated for inference training, validation, etc., we supply *Relate* with the mean value of our simulated distribution.

### Overview of Graph Convolutional Networks (GCNs)

Graph convolution (Kipf and Welling 2017) works by aggregating information at each node in a graph by combining data from its neighbors (those nodes with which it shares an edge). The input to a graph convolution operator is a graph or a set of nodes and edges (*N*, *E*), where each x ∈ N is some n-dimensional vector containing information about a given node, and each element of *E* is a pair of node indices representing a connection between those two nodes. The edges of a graph are often specified instead by an adjacency matrix ***A*** which specifies the topology of the graph: i.e. ***A*** has a value of 1 at entry *ij* if node *i* and node *j* share an edge and a value of 0 otherwise. Formally the graph convolution can be written in matrix form as:

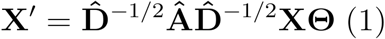

where **X** is a matrix consisting of the *n* node feature vectors 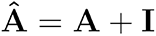 as the identity matrix and 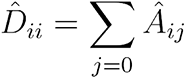 as the diagonal degree matrix. Here features are first aggregated according to the connectivity specified in **A** including the features belonging to the node itself (hence the addition of the identity matrix specifying self-connectivity, a “self-loop”) and then normalized using the degree matrix to avoid exploding gradients. Then, they are transformed according to some learnable matrix Θ optimized through gradient descent during training. For clarity, the node-wise formula is:

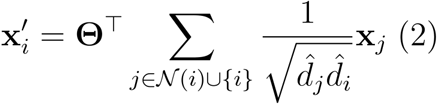

where the degree of node (including self-connectivity) *i* is 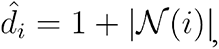, with 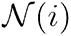 being the set of neighbors of node *i*, and *x_i_* being the node features for node *i*. Thus, in each graph convolution step, each node’s features are updated to the weighted sum of all its neighbors features including its own (i.e. the features of the set of nodes 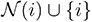, as specified in the summation). Specifically, each contribution of each node *j*’s feature vector is weighted by the reciprocal of the square root of the product of node *j*’s degree and node *i*’s degree and the degree of its neighbor, as in the denominator of Equation 2. Then, that resulting sum is transformed according to the learnable matrix Θ. This is called a “convolution” on graphs as it is analogous to convolutions on images where the “kernel” of a graph convolution operates over each local neighborhood of each node much like an image convolution kernel operates on some rectangular region around each pixel (see example in Figure S4).

In the case of genealogical trees, the node features returned by each graph convolution operation are a function of those of its parent and its two children if it is an internal node, its two children only if it is the root node, and its parent only if it is a leaf node. For our input graphs we used the following features: the age of a node (the time in log generations that the coalescence event associated with it occurred mean and standard deviation normalized, or 0 for leaf nodes), the number of mutations on the branch from its parent (mean and standard deviation normalized via the training data), and a one-hot encoded label vector specifying whether the node is internal (i.e. ancestral) or external (present day) respectively, and the population label if the node is external (for internal nodes the population is considered unknown). Thus, if there are *k* populations the size of the label vector is *k* + 1. Note that the definition of graph convolution allows for directionality of the edges, i.e. one can specify whether node *i* passes information to node *j* in each graph convolution operation or vice-versa. Here we specify that the edges are unidirectional with the direction being child to parent. Thus, for our GCN 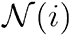 is the set of child nodes of node *i*., and no update is performed for external/leaf nodes in the tree 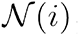 is the empty set, and thus for those nodes 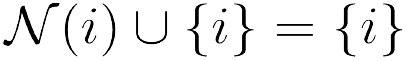, meaning only the focal node is considered during the convolution.

### GCN architecture and training

The GCN architecture implemented here (illustrated in Figure 1) consists of two components: a graph-learning network, or layers that operate on graphs, and two recurrent networks that operate on sequences of feature vectors. This GCN architecture contains elements specifically designed to the task of learning from tree sequences and is meant to be an advancement of the use of GCNs in population genetics, which is why it is not an “out of the box” solution such as the CNN described below. First, each tree in the input sequence has its node features transformed via a learned affine transform with bias, to which the original features are concatenated to via a “skip connection” before each tree is then input separately to a series of graph convolution layers with skip connections, comprising a graph convolution “block” of layers. At each layer, the output from the previous layer is transformed via convolution and added to itself, then the result is normalized and passed to the ReLU activation function (Agarap 2018). For normalization in the GCN layers we use LayerNorm (Ba et al. 2016). The graph convolution layer we chose for our architecture is the Graph Attention Convolution (GATConv) from (Brody et al. 2022) implemented in torch-geometric (Fey and Lenssen 2019; Paszke et al. 2019). It is a modified form of the proposed convolution operator that we detail in the previous section in that node features from each neighbor are linearly transformed *before* they are summed in the update step and are weighted by learned attention coefficients instead of simply the degree of the nodes, enabling the network to potentially learn which neighbor’s features are more or less relevant for accomplishing the given regression task. In short, weighting by node degrees as described in the previous section prevents gradient explosion whereas weighting by attention prevents gradient explosion *and* has the benefit of learning the most optimally-informative connections in the graph for a given task (Brody et al. 2022). Formally the convolution with attention is specified as:

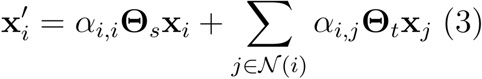

Where the attention coefficients *α* are computed as:

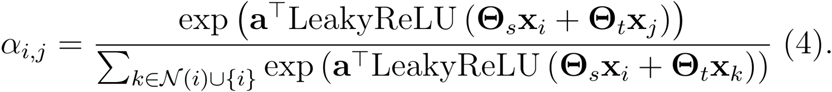

With 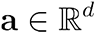 and 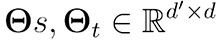 being learnable parameters weighted during training where *d* and *d′* are the dimension of the input features and output features respectively and *R^d^* being a *d*- dimensional vector of real numbers. In each GATConv layer, attention coefficients *α_ij_* are computed which selectively filter messages from neighboring nodes. This attention mechanism is comprised of the same two learnable affine transformation matrices used in eq. 5, Θ*_s_* and Θ*_t_*, which transform the focal node and its neighbor’s node features respectively, and **a** being a trainable weight vector onto which the summed features are projected via the dot product and then normalized over all neighbors to sum to one using the softmax function. In this way, the attention mechanism hopes to weight the messages from neighboring nodes with *α_ij_* such that returned features have the most impact on loss reduction for a given target. Note that self-attention coefficients expressing how much the focal node’s own features contribute to the features in the next layer (*α_ij_*) are also computed in a similar manner.

After passing through the GATConv layers, features obtained from the nodes in each tree are separately passed through a Gated Recurrent Unit (GRU; (Cho et al. 2014)) and the hidden features are returned as the compressed vector representation of each tree in the sequence. For each GRU’s input, the order of nodes in the tree is as follows: the leaf nodes are first in the sequence, appearing in the same order as they do in the sample returned by the simulator, and are followed by the internal nodes, which are ordered by the time in generations that they appear in the tree. Although the order of these nodes is therefore somewhat arbitrary, we reasoned that using a GRU could potentially provide two advantages over a simple dense layer that does not assume any sequential order of its inputs 1) the GRU may learn to take advantage of any consistency in the sequential order of nodes as written out by *Relate*, and, more importantly, 2) using a recurrent neural network layer like a GRU allows us to use a relatively small number parameters, while still having a sizeable number of weights per input node (because the same set of weights are shared across input nodes).

The vector emitted by each GRU is concatenated to a “tree summary vector” containing the following features about the original tree: the time of coalescence of all individuals in the tree, the mean, median, and standard deviation of coalescence times, the mean, median, standard deviation, skew, and maximum of the branch lengths in the tree, the midpoint of the chromosomal segment the tree corresponds to (scaled from 0 to 1, with 0 being the left end of the simulated chromosome and 1 being the right end), and the fraction of the entire sequence accounted for by this segment. The resulting vectors, one for each input tree, are then concatenated into a single matrix, which is in turn passed through another GRU that emits a feature vector summarizing the entire tree sequence. A global vector (the “tree sequence summary vector”) which consists of the mean, standard deviation, and median of the tree summary vectors, as well as the number of graphs in the original sequence, is embedded using an affine transform with bias and then concatenated to the resulting hidden vector from the previous GRU, resulting in a 1D vector containing information extracted from the tree sequence as a whole. Each global vector and pre-computed tree summary vector for the graphs is mean and standard deviation normalized via the training data set in preprocessing. Finally, this is passed through three fully connected layers (a multi-layer perceptron, “MLP”) with the first two using ReLU activations and batch normalization (Ioffe and Szegedy 2015), and the final (output) layer using no activation function. This results in classification scores which could be interpreted as posterior probabilities by applying the softmax function, or in the case of regression, *z*-scores for the various values to be predicted (all *y* values for the regression tasks are standardized via the mean and standard deviation computed from the training set). We provide a table specifying the dimensionality of the inputs and outputs in each layer of our network as well as the activations used for clarity (Table 5). We note that the chosen architecture was the result of some trial and error in which different types of graph convolution as well as serial learning networks, such as 1D convolution in the place of GRUs, were attempted, and the architecture we present here is the result of that process. While we did not conduct a systematic exploration of different architectures and hyperparameters to find an optimal design, we did experiment with removing parts of this architecture and altering neural network depth to assess the impact of some of our architectural choices (see **The impact of neural network depth on performance** and **The impact of removing GCN architecture components** in the Results). Nonetheless, there may be other architectures that would perform better on the set of population genetic benchmarking tasks considered here. In addition, this set of benchmarking tasks was by no means exhaustive, and while we found our choice to generalize well to these problems, we cannot be certain that our architecture would be well suited for all possible tasks.

When passing the ragged sequences to the final GRU layer they are each padded to the maximum length observed in the given batch. For all the experiments however, we first downsample the tree sequences if they are > 128 trees in length by randomly selecting a sequential window of 128 trees and omitting the others. This cap on tree sequence lengths allowed us to increase the batch size while still being able to train using a single GPU. For some tasks, namely in the case of recombination rate and demographic parameter regression, this is far more than the average number of trees observed in the chosen window size and thus most tree sequences are passed in full to the network. However, in the case of the sweep detection problem, there are often far more than 128 trees in the tree sequence, and thus many of the replicates passed to the network are downsampled sequences. Further details of how we downsample the tree sequences and the average amount of downsampling are given in the sections detailing each task.

For classification tasks, we used categorical cross entropy as our loss function to be minimized during training:

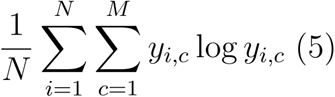

Where *y_i,c_* is a binary indicator of the correct class and *y_i,c_* is our posterior probability estimate that the observation came from this class where *i* is the replicate index, *C* is the class index, *M* is the number of classes, and *N* is the batch size or number of replicates considered in each gradient step. For regression tasks, we use a variant of L1 loss (i.e. mean absolute error (MAE)) because of L1’s relative insensitivity to outliers; specifically, we used the “smooth L1” loss function, whose slope decays at values closer to zero (Girshick 2015). However, note that for all methods and tasks we report final regression accuracies in root mean-squared error (RMSE) to better compare to Flagel et al. (2019). *y* values for the regression tasks are standardized via the mean and standard deviation computed from the training set. We trained using the ADAM optimizer (Kingma and Ba 2017) with a learning rate of 1e-5 for up to 100 epochs or until their validation accuracy stopped increasing for 10 consecutive epochs. For each problem we specify the batch size in the sections below.

Although at surface level our approach appears to be similar to the GNN used by Korfmann *et al*. (2023b) GNN*coal* method, there are several notable differences. For example, while GNN*coal* represents each tree as an undirected graph, we used directed graphs with edges pointing up the tree from child node to parent node. GNN*coal* also uses a hierarchical graph- pooling strategy to compress each tree down to a graph with a smaller number of nodes, while we performed no such pooling operation and instead embed the information extracted from each tree with a learnable GRU, later using a second GRU to extract information from the sequence of trees. Finally, our graph convolution operator features learnable attention coefficients, not present in GNN*coal*. Although it is possible that the GNN*coal* architecture could be adapted to perform well on the tasks we examine in this paper, we did not investigate this possibility as our primary focus was to determine whether a neural network trained to examine inferred tree sequences could perform as well as one directly examining the raw genotype matrix—as we show in the Results below, the GCN presented here was sufficient for this purpose.

Because input tree sequences are of variable length, here we define the dimensionality of a single batch as (*B*, *T*×*N*) where B is the batch size (number of tree sequences in a batch) and *T*×*N* is the number of trees × the number of nodes. For each node, we calculate a set of *F* features, giving us an input shape of (*B*, *T*×*N*, *F*) as shown in the “input shape” column for the first layer Table 5 (the “node embedding” row). The inputs to the graph convolution layers are a 2-D tensor of shape (*T*×*N*, 26+*F*), where *T*×*N* is again the total number of nodes in the tree sequence, and 26 features per node resulting from concatenated with the original node features from the first skip connection), and a list of edge indices of shape (2, *E*) specifying the pairwise connections between nodes where each row is a pair of node indices the edge connects, similar to an adjacency matrix this information is used by the graph convolution layers to utilize the node connectedness. The input and output shapes of the skip connection layer, reshaping layer, and first GRU layer that follow the graph convolution layers are also determined by *T*, *N*, and *F* as shown in Table 5. When considering the input to the second GRU layer, *T*’ refers to the padded tree sequence length, which is the maximum observed number of trees among the *B* tree sequences in a given batch. This padding is performed because all tree sequences in a given batch must have the same length, and we chose to increase the length of tree sequences to T’ by concatenating tensors filled with zeros to the end of the GRU’s input tensor rather than downsampling the tree sequence by removing trees. Note that this padding was done using PyTorch’s `pack_padded_sequence` function that allows for more efficient computation by allowing *T*’ to vary across batches rather than setting it to the maximum length across all batches and instructing the GRUs to ignore the padded part of the sequence during forward and backward computations (Paszke et al. 2019). This is followed by another concatenation layer and then several fully connected layers culminating in the output layer. The input and output shapes for all of these are shown in Table 5. All benchmark tasks except the introgression-detection task use identical output shapes for all layers except the final output layer, which is task-dependent. In total, the chosen architecture has 228,837 parameters for the demography and positive selection problems, 229,771 parameters for the introgression classification problem, and 228,381 parameters for the recombination rate problem.

### CNN training

The CNN used for comparison was the Pytorch (Paszke et al. 2019) implementation of the ResNet34 architecture (He et al. 2015). ResNets are a type of CNN that feature skip or residual connections that merge output from the convolutions with the output from previous layers via addition, and have been shown to perform competitively on traditional image recognition tasks (He et al. 2015; Reddi et al. 2020). Models were trained for up to 100 epochs or until their validation accuracy stopped increasing for 10 consecutive epochs. For classification tasks (selection detection, introgression detection) categorical cross entropy loss was used, while smooth L1 (Girshick 2015) loss was used for regression tasks (rho estimation, demographic inference). The ADAM optimizer (Kingma and Ba 2017) with default settings (lr=0.001, betas=(0.9, 0.999)) was used for all networks. All genotype matrix inputs were formatted to be of shape (populations, individuals, polymorphisms) and were padded with zeros to the max observed number of polymorphisms in each case. The rows were sorted based on cosine distance using seriation as described in (Ray et al. 2023) using Googles OR-tools package (Perron and Furnon 2019); note that the sorting algorithm and similarity metric differ from that used by (Flagel et al. 2019) as we found the seriation approach to perform better on the task of identifying introgressed haplotypes (Ray et al. 2023). Data was represented as binary encoding of 0 or 1 (ancestral or derived) for all tasks.

The batch size for the CNN for both regression cases was 32. The batch size for the selection problem was 10 as the padded size was very large at 5000 SNPs. The batch size for the introgression problem was 96. The CNN used for comparison was the ResNet34 architecture (He et al. 2015) which was implemented in PyTorch. ResNets are a type of CNN that feature skip or residual connections that merge output from the convolutions with the output from previous layers via addition, and have been shown to perform competitively on traditional image recognition tasks.

### Test Cases

#### Inferring the historical recombination rate

Simulations for training and benchmarking performance on recombination rate inference were done using *ms* (Hudson 2002). In particular, for each simulation replicate a population size *N* was drawn from the set {1000, 2000, 5000, 10,000, 20,000, 50,000}, and, after setting the locus size to *L*=20kb and the mutation rate to *μ*=1.5×10^-8^, a recombination rate *ρ*=4*NrL* was chosen such that the per-base pair crossover rate, *r*, was drawn from a truncated exponential distribution with lower and upper boundaries of 10^-8^ and 10^-6^, respectively. The sample size was set to 50 haploid chromosomes. Simulation and formatting resulted in 237,004 replicates for training the CNN, and 12,104 for its validation.

Recombination rates values were log-scaled and then standardized to a *z*-score using the training set mean and standard deviation. Note that the validation and test data were also standardized using the training set’s mean and standard deviation to prevent data leakage from the training to the validation/test data. Because the number of segregating sites differed across simulation replicates, the data was padded to 413 polymorphisms or the maximum observed number of polymorphisms in the simulated replicates. The median number of trees returned by *Relate* (computed from training data) was 13 with a standard deviation of 24.05.For the GCN, the training and validation sets had 231,208 and 11,791 replicates respectively.The test set used to compare the CNN and GCN had 4,866 replicates.

#### Detecting selective sweeps

We benchmarked accuracy on the task of detecting selective sweeps by adopting the approach of (Schrider and Kern 2016; Flagel et al. 2019) in that we define the problem as a classification task with 5 classes: hard sweeps (a sweep of a *de novo* mutation), hard-linked regions (i.e. those close to a hard sweep), soft sweeps (selection on standing variation), soft-linked regions (close to a soft sweep), and neutrally evolving regions. Sweeps were simulated using *discoal* (Kern and Schrider 2016) with the same simulation parameters used by Schrider and Kern (Schrider and Kern 2017) to generate training and test data for the JPT sample (Japanese individuals from Tokyo) from Phase 3 of the 1000 Genomes dataset (Auton et al. 2015).

For the hard and soft sweep classes, the target of selection was located at the center of the simulated chromosome. For the sweep-linked classes the target of selection was located outside of the central “sub-window” (which accounts for 1/11 of the total length of the selected chromosome). For this task we consider a simulated example to be a sweep if the central polymorphism in the sampled window is the target of selection. Sweep-linked classes are defined as having a sweeping allele in the sampled window but not in the center, while the neutral class does not have any selection in the simulation. For sweep scenarios we condition on the sweeping allele having reached fixation at time *τ* ∼ *U*(0, 2000) generations ago. The sample size was set to 104 haploid individuals. Genotype matrices were either padded or cropped to 5000 polymorphisms. The training and validation sets for the CNN had 236,232 and 26,432 examples, respectively.

For the tree sequences, sequences containing more than 128 trees were downsampled as in the other problems, but instead of sequential window sampling we sampled trees by setting the probability that a tree was sampled by the fraction of the chromosome that it spanned, and then sampling with replacement while keeping the original ordering intact. This was done because a large proportion of simulations had >> 128 trees in sequence (the median length across the five categories was 107 with standard deviation of 111.2) and non-linked replicates can have selection almost anywhere along the simulated chromosome, thus contiguous window sampling could often miss the relevant signal. The training and validation sets for the GCN had 224,555 and 25,445. The test set used to compare the performance of the CNN and GCN had 22,480 replicates. The training, validation and testing data were evenly balanced across the five categories.

#### Detecting introgressed loci

We simulated data under the same *Drosophila* demographic model from Ray *et al*. (2023) as data for a 3-class classifier problem to benchmark each networks’ ability to detect the presence and direction of introgression within a genomic region. In brief, we simulated a pulse migration event from *D. simulans* to *D. sechellia* at a random time sampled from *U*(0.3, 0.5) × *T*, where *T* is the population split time inferred by bootstrapped ∂a∂i (Gutenkunst et al. 2010) replicate inferences on *D. simulans* inbred and *D. sechellia* wild-caught genomes. In contrast to the parameterization simulated by Ray et al. (2023) which used more recent introgression events, we chose the introgression time distribution to be *U*(0.3, 0.5) × *T* to make the problem a more difficult benchmark (because older introgression events have introgressed haplotypes that are less diverged between the donor and recipient population, and also that have had more time to be broken up by recombination events after the introgression event).

For this task genotype matrices are formatted in the shape of (*samples*, *populations*, *individuals*, *polymorphisms*). Data was padded to 665 polymorphisms or the maximum observed number of polymorphisms in simulated replicates. The median number of trees returned by *Relate* (computed from training data) was 68 with a standard deviation of 11.56.For the GCN, the training and validation sets had 232,050 and 25,950 replicates respectively. For the CNN, the training and validation sets had 244,740 and 13,260 replicates respectively. The test set used to compare the methods contained 12,900 replicates, and the training, validation, and test sets were all class-balanced.

#### Demographic inference

As in (Flagel et al. 2019) we tested the ability of our networks to perform demographic inference by simulating a 3-epoch demographic model with 5 parameters: the present-day population size (*N*0 which was drawn from *U*(100, 4×10^4^)), the time of the most recent population size change (*T*1 ∼*U*(100, 3500)), the population size during the middle epoch (*N*1 ∼*U*(100, 5000)), the time of the ancient population size change (*T*2, which was set to *T*1 plus a value drawn from *U*(1, 3500)), and the ancestral population size (*N*2 ∼*U*(100, 2×10^4^)). These data were simulated using *ms* and the corresponding scripts can be found at https://github.com/flag0010/pop_gen_cnn/tree/master/demography.For the CNN, the training / validation had 94,036 and 5,164 replicates respectively.

The data was padded to 652 polymorphisms or the maximum observed number of polymorphisms in the simulated replicates. Target values were log-scaled and standardized prior to training and testing, with the validation and testing data being standardized according to the mean and standard deviation of the log-scaled training data. For the GCN, the training and validation sets had 82,820 and 9,168 replicates respectively. The median number of trees in sequence was 53 with a standard deviation of 37.0. The test set used to compare the two methods contained 3,995 examples.

## DATA AND CODE AVAILABILITY

All code associated with this work can be found at https://github.com/SchriderLab/TreeSeqPopGenInference. Model weights can be downloaded from https://drive.google.com/drive/folders/1WRf8pecfRavOQjmZF7Otr4lXbcouVunm.

## ACKNOWLEDGMENTS

The authors thank Will Booker for comments on the manuscript, and members of the Kern and Ralph Labs for feedback on preliminary data. LSW was supported by NIH award R01AI153523, DDR was supported by NIH award R01HG010774, and DRS was supported by NIH award R35GM138286.

