## Supplementary Material for "Tree sequences as a general-purpose tool for population genetic inference"

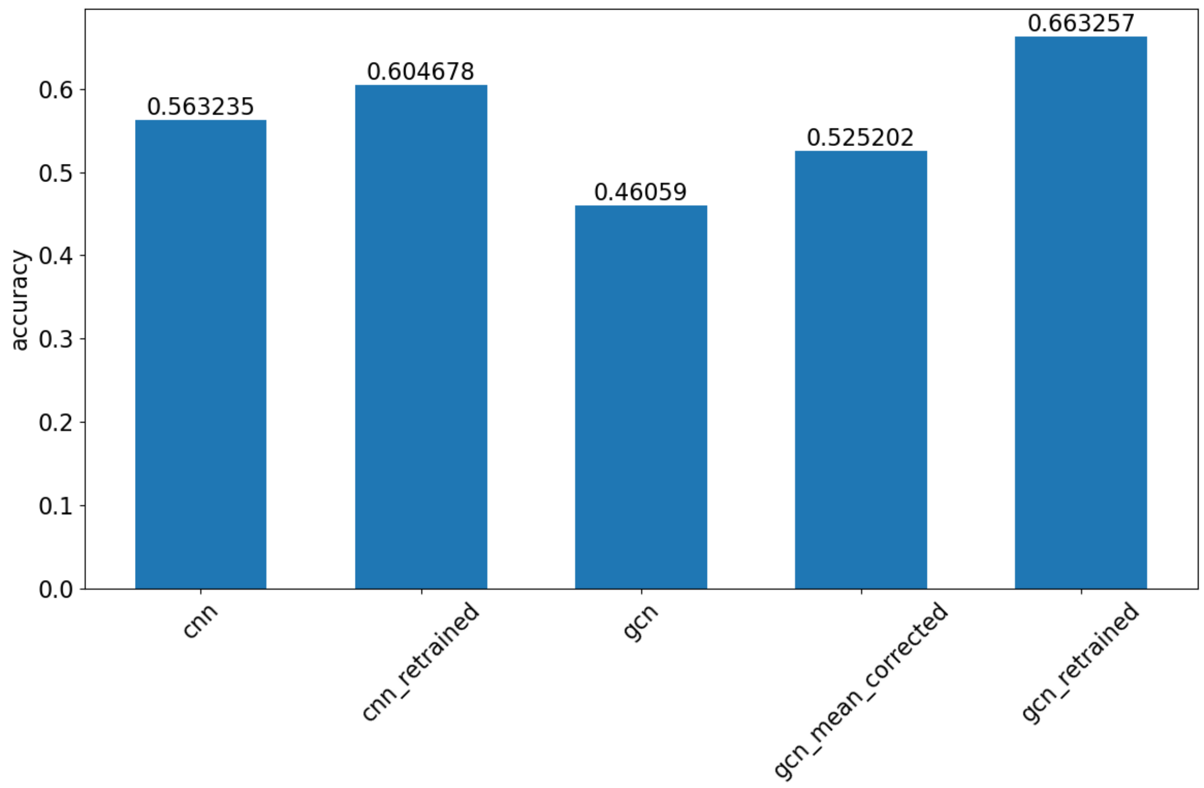


Figure S1: Accuracy on a common evaluation set for the selection window classification problem where for each replicate $\theta$ and $\rho$ are perturbed identically and independently away from the original values by a multiplicative factor drawn uniformly from 0.5 to 1.5. The “retrained” bars show the accuracy after retraining the network architecture with a portion of the training data now simulated in the same way. For the GCN, we also computed the accuracy when we simply correct the mean and standard deviation of the input tree and tree sequence feature vectors to account for the new distribution.

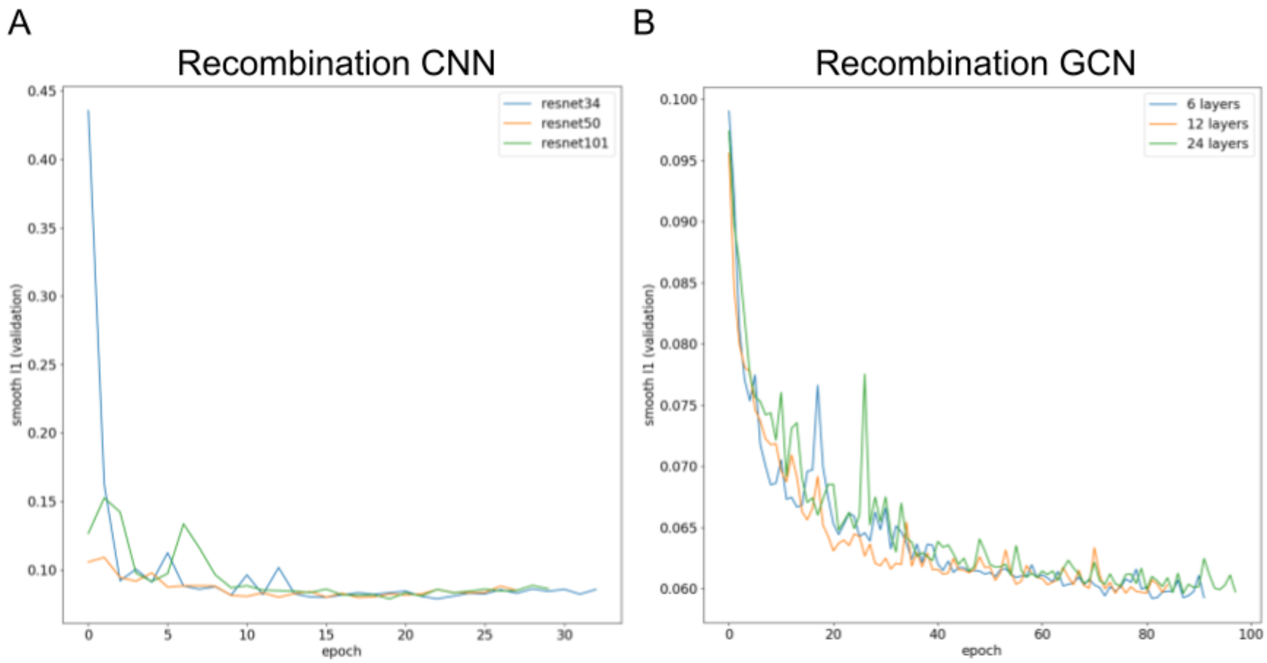


Figure S2: Training curves for predicting recombination rate for different choices of depth for the (**A**) ResNet CNN architecture and (**B**) GCN architecture. Accuracy across training epochs is measured by smooth L1 loss (or mean absolute error, MAE) over epochs on held-out validation data each epoch. The ResNet34 and 6-layer architectures were the ones we used for our main CNN and GCN results, respectively.

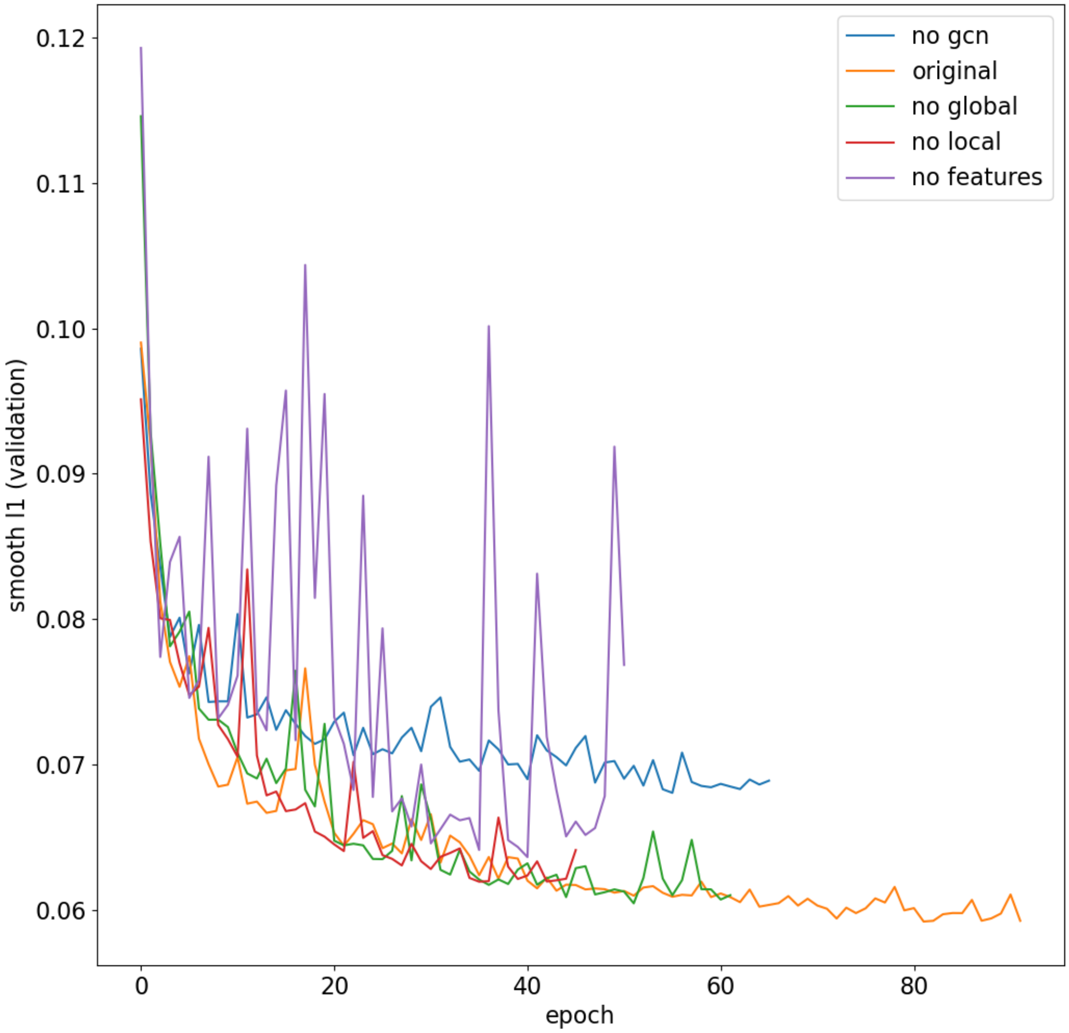


Figure S3. The validation loss trajectories observed on the recombination estimation task when removing different parts of our GCN architecture and associated inputs. Original: the full GCN architecture and input; no GCN: the graph convolution layers have been removed, so the only information included are the tree summary features and tree-sequence summary features described in the Methods; no global: the tree sequence features are omitted, and the graph convolutions and tree features are included; no local: the tree features are omitted, and the graph convolutions and tree sequence features are included; no features: the tree and tree sequence features are omitted, so only those features extracted by the graph convolutions are used.

A

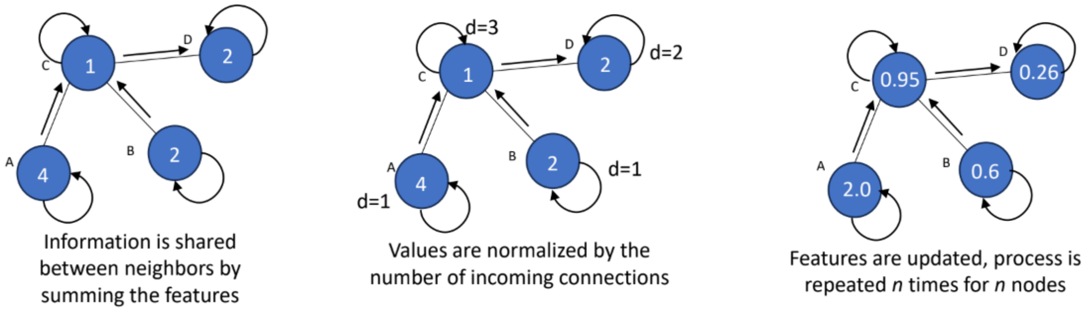


B

| Node | Weight | Input Features | Degree | Calculation | Result $(x'_i)$ |
|---|---|---|---|---|---|
| A | 0.5 | 4 | 2 | $(\frac{1}{\sqrt{1}*\sqrt{1}}*4)*0.5$ | 2.00 |
| B | 0.3 | 2 | 2 | $(\frac{1}{\sqrt{1}*\sqrt{1}}*2)*0.3$ | 0.60 |
| C | 0.25 | 1, 4, 2 | 4 | $((\frac{1}{\sqrt{1}*\sqrt{3}}*4)+(\frac{1}{\sqrt{1}*\sqrt{3}}*2)+(\frac{1}{\sqrt{3}*\sqrt{3}}*1))*0.25$ | 0.95 |
| D | 0.1 | 2, 1 | 2 | $((\frac{1}{\sqrt{3}*\sqrt{1}}*1)+(\frac{1}{\sqrt{1}*\sqrt{1}}*2))*0.1$ | 0.26 |

Figure S4. (A) Diagram of graph convolution and (B) resulting parameters and calculations to demonstrate graph convolution. In this example features are simplified to a vector containing a single learnable parameter, denoted by “weight”, and a single scalar value for initial input features.

Table S1. Simulation and sampling parameterizations for each benchmark task.

| Task | Selection | Introgression | Recombination | Demographic inference |
|---|---|---|---|---|
| *N* (Population size(s)) | Population sizes follow the shape of the PSMC model estimated by Auton et al. (2015) for the JPT human population. The present-day population size is set to ~2.29e4. | Mean values:<br>$\bar{N}_{anc}$ = 2.34e5<br>$\bar{N}_{sech}$ (*sechellia* size post-split) = 4.99e6<br>$\bar{N}_{sim}$ (*simulans* size post-split) = 1.74e4<br>$\bar{N}_{sech\text{-}0}$ (*sechellia* present-day size) = 6.42e4<br>$\bar{N}_{sim\text{-}0}$ (*simulans* present-day size) = 2.23e6<br><br>See additional notes for more information. | Constant *N* for each replicate (drawn from {1000, 2000, 5000, 1e4, 2e4, 5e4}) | Three epochs:<br>$N_0$ (present-day): ~$U(100, 4\times10^4)$;<br><br>$N_0$ (middle epoch): ~~$U(100, 5000)$;<br><br>$N_2$ (oldest epoch): ~$U(100, 2\times10^4)$ |
| Time of population splits/size changes | The timing of size changes also follow the JPT PSMC model from Auton et al. (2015). | Mean value of *simulans-sechellia* split: 6.29e4 years ago (assuming 15 gen/year). Ancestral size is constant. *D. simulans* and *D. sechellia* populations experience continuous exponential size change post-split. | N/A | Size change times:<br>$T_1$ ~$U(100, 3500)$<br>$T_2 = T_1$ + ~$U(1, 3500)$ |
| Additional notes on demographic model | All past population sizes are set relative to the present-day population size, which was selected as described in Schrider and Kern (2017). | A variant of the *simulans-sechellia* model from Ray et al. (2015), with the timing of introgression changed to ~$U(0.3, 0.5) \times T$, where *T* is | Constant-sized demographic model where each rep has a different population size and recombination rate. | Piecewise-constant 3-epoch model |

| | | | | |
|---|---|---|---|---|
| | | the split time. | | |
| $\mu$ (mutation rate) | ~$U$(2.18e-9, 2.18e-8) | 5e-9 | 1.5e-8 | 1.2e-9 |
| $r$ (recombination rate, measured in crossovers per base pair per generation) | Drawn from a bounded exponential with mean of 1e-8, with rejection sampling such that only values <3e-8 are accepted. | Drawn uniformly from 1.67e-8 to 5e-8. | Drawn from an exponential with mean 2.15e-7, with rejection sampling such that only values between 1e-8 to 1e-6 are accepted. | 1e-8 |
| sample size | 104 | 34 | 50 | 50 |
| $L$ (sequence length) | 110,000 | 10,000 | 20,000 | 1,500,000 |
| CNN Training set size | 236,232 | 244,740 | 237,004 | 94,036 |
| CNN Validation set size | 26,432 | 13,260 | 12,104 | 5,164 |
| GCN Training set size | 224,555 | 232,050 | 231,208 | 82,820 |
| GCN Validation set size | 25,445 | 25,950 | 11,791 | 9,168 |
| Test set size | 22,480 | 12,900 | 4,866 | 3,995 |
